# The capacity for correlated semantic memories in the cortex

**DOI:** 10.1101/321703

**Authors:** Vezha Boboeva, Romain Brasselet, Alessandro Treves

## 1 Introduction

One of the most fascinating aspects of the human brain is its ability to ascribe significance to and recognize meaning in objects and events, and more generally to make sense of the world. Semantic memory, comprising our acquired knowledge about the world, can be imagined to reflect, in its statistical structure, the complex, distributed, policentric structure of the neocortex where it resides. In contrast, the relatively much simpler network structure of the hippocampus, in particular of its CA3 field, where episodic memories have long been thought to be at least initially represented by unique patterns of neural activity, may lead to the limited set of outcomes of episodic memory retrieval: either the pattern is retrieved, or not. In the first case, retrieval, subjects *remember* what happened in the episode, in the second they do not, although they may still *know* many of the elements in the episode, likely as they reconstruct them with input from semantic memory. This is the basis for *remember/know* paradigms [1] that assess hippocampal contribution to memory retrieval. But how can the statistical structure of the memory representations themselves be characterized?

### 1.1 Correlations

In the case of episodic representations in the hippocampus, one straightforward hypothesis about their statistics is that they are largely uncorrelated: each representation is set up, e.g. in CA3, independently of other representations already stored there, under the influence of the Dentate Gyrus [2]. Then the representations are roughly at the same distance from each other in activity space, i.e. they are *ametric:* relations of being closer or farther away, or in the middle between another pair, lose their meaning. This may seem at odds with the best studied neural representations in the hippocampus, spatial representations in rodents, which reflect the continuity of space, where being close or distant is clearly defined. As soon as we move, however, from the representation of different locations in the same restricted spatial context to the representation of different contexts, the phenomenon of global remapping suggests that the notion of ametric representations is relevant. Indeed, it has been observed that even very similar spatial contexts are represented in rat CA3 by completely different, essentially uncorrelated representations [3]. Correspondingly, a measure of metric content has been shown to “increase” in human subjects who can rely *less,* due to incipient Alzheimer, on their hippocampal representations [4]. If hippocampal representations can be said to be ametric, what is the nature of the metricity observed, by contrast, in semantic representations in the neocortex?

Direct access to individual semantic representations through single unit recordings is of course very limited, and not just in the human brain, because of their very distributed nature. Multivoxel pattern analyses from fMRI are consistent with a complex web of correlations [5, 6], but their resolution is limited and so is the characterization of the statistical properties of those correlations. A simple alternative, however, is to assess the nature of the correlations among the semantic items themselves, rather than probing their representations in the brain. This can be done by utilizing any of a number of databases, where a set of semantic items have been described in terms of the features or attributes people associate with them. As a simple toy example, we took the *p* = 60 nouns used in a recent fMRI study [7]. We computed the pairwise correlation between these nouns, as measured by a set of intermediate or surrogate features, such as the co-occurrence with a set of verbs within a sentence in the corpus.

In Fig. 1(a), we report the correlation matrix of the nouns, from which it can be seen that the pattern of correlations cannot be described by any simple schema. One way of thinking about this organization, that is ubiquitous in the semantic literature, is to think of concepts organized in a hierarchy, or a tree [8, 9]. Such models, in their descriptive and generative formulations, are dramatic oversimplifications that ignore important features of the data, such as the prevalence of concepts intermediate between other concepts (see Sect. 2.2.1). As an example, in Fig. 1(c), we report the correlation that an extreme hierarchical model would see in such data. Unsurprisingly, it is made of clusters with large within correlation and no between correlation. It is apparent, from the comparison with Fig. 1(a), that this hierarchical model fails to represent off-block values with high correlation.

**Figure 1:**
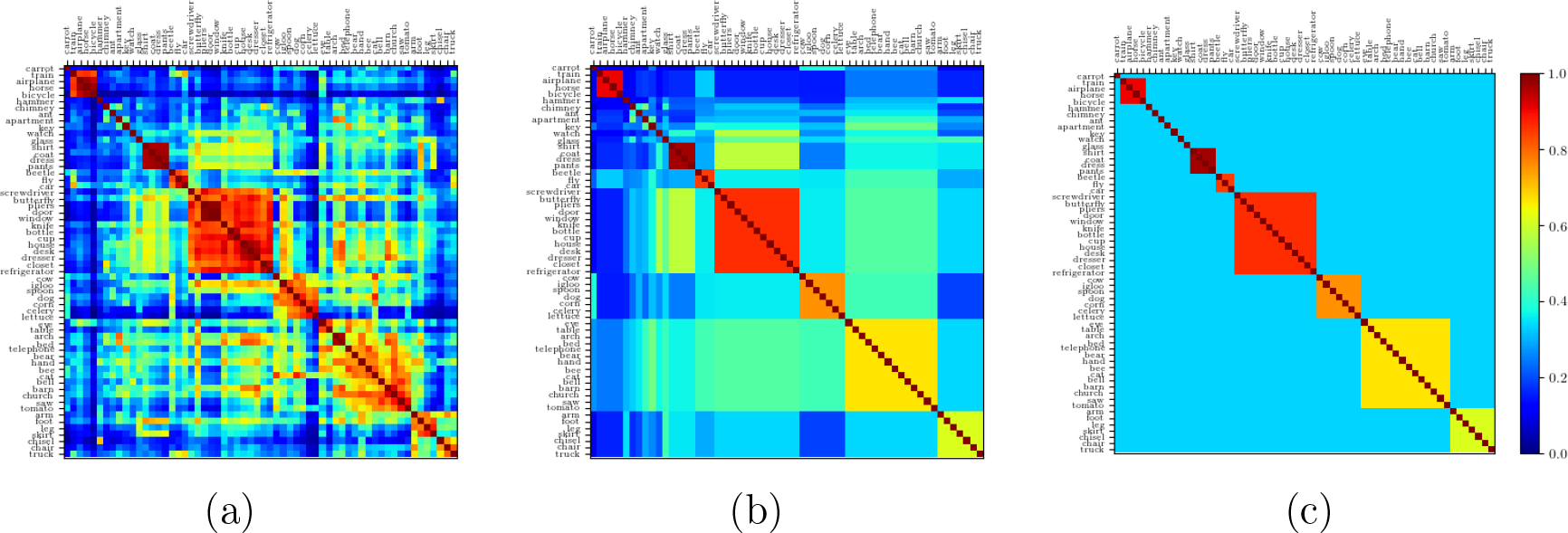
(**a**) Original correlation matrix. (**b**) Correlation matrix obtained by replacing each within-block entry with the mean correlation value of that noun cluster and each off-block entry with the mean correlation between clusters. The clusters are obtained through the application of a standard clustering algorithm to the original correlation matrix, Fig. 1(a). (**c**) Strictly ultrametric correlation matrix obtained by again replacing each within-block entry with the mean value within that block, and now each off-block entry with the overall off-block mean.

A less dramatic simplification consists in considering that the correlations between individual concepts belonging to the distinct clusters can be well approximated by the mean correlation between clusters. Such a simplification yields Fig. 1(b). To quantify the validity of this simplification, one can measure to what extent the distance relations between the concepts match the fully hierarchical limit case [10]. This index, called the ultrametric content (see App. C) can be computed once correlations are translated into a measure of distance (Fig. 2). The “soft” hierarchical structure of the matrix in Fig. 1(b) yields an ultrametric content index of 0.61, to be compared with the value 0.5 for the original data. The fully hierarchical matrix, Fig. 1(a), has an ultrametric content of 1. On the other hand, ametric, independently generated representations, as observed in CA3, have an ultrametric content close to 0. Semantic relations, we can conclude, are complex, and the ultrametric content of 0.5 is as far from the purely ultrametric as from the trivial ametric limit.

**Figure 2:**
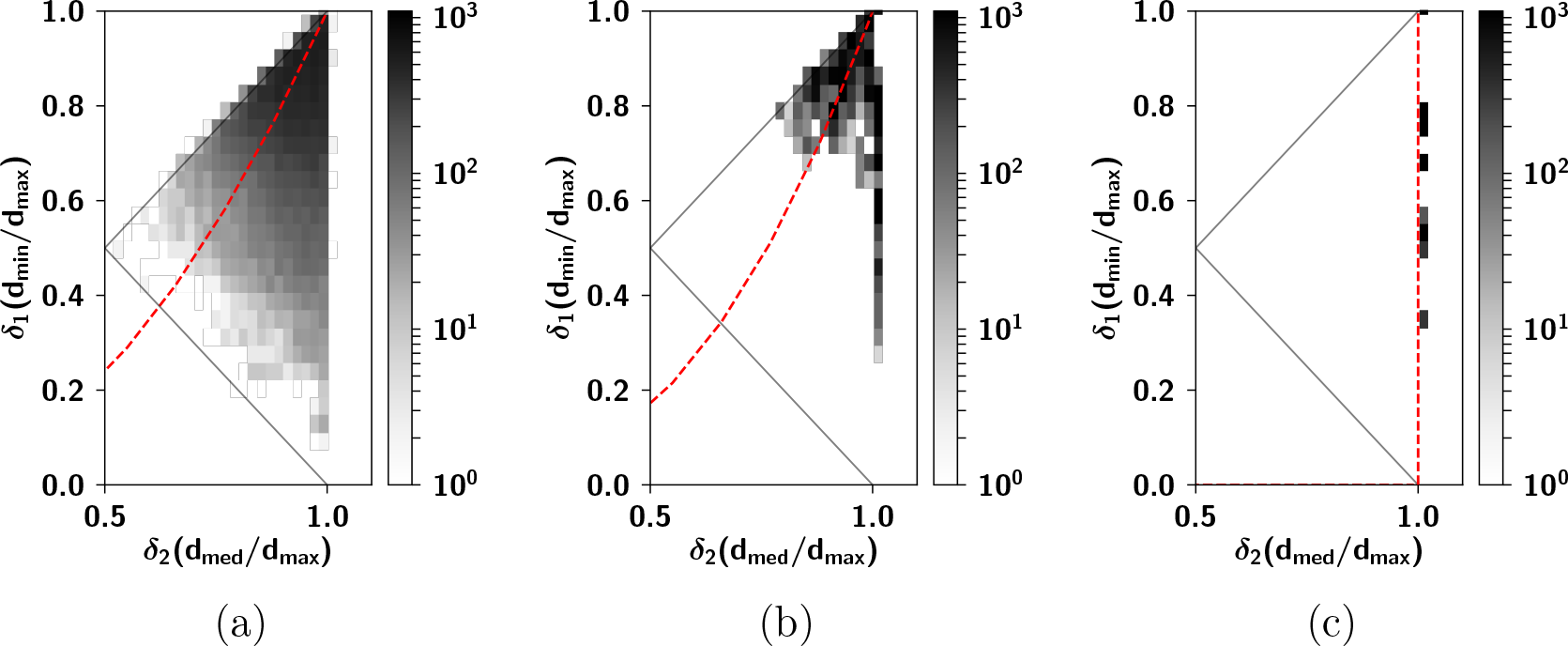
(**a**) Two-dimensional logarithmic density plot of the ratio between the intermediate and the longest edge vs. the ratio between the shortest and the longest edge, in the triangles created by extracting quasi-distances for the triplets of nouns taken from Fig. 1(a). The triplets are scattered, with an ultrametric content of 0.5. (**b**) Same as Fig. 2(a), with triplets taken from Fig. 1(b). The triplets have less scatter and yield an ultrametric content of 0.61. (c) Same as Fig. 2(a), with triplets taken from Fig. 1(c). Here triplets constitute isosceles triangles with two long sides, as can be seen from the alignment of the ratios along the vertical line **d*_*med*_ = *d*_*max*_.* The ultrametric content (see App. C) is exactly 1. In all three panels, the dashed red line corresponds to the line of constant ultrametric content index.

However, it is the statistical independence of memory patterns that had made available most of the mathematically sophisticated analyses of autoassociative networks. While these analyses have been successfully used to describe the CA3 circuit, it does not seem like they can be applied to semantic memory, which has in the shared structure between memories its *raison d’être*. Still, in exploring variants of the Hopfield model, which initially featured uncorrelated patterns, the challenge of storing correlated patterns was eventually taken up. One of the earliest attempts to introduce correlations was through an algorithm that arranged patterns on a tree [11, 12], in which the upper nodes correspond to classes of items and the lower nodes, each branching from a single upper node, correspond to exemplars of a class. Subsequently it was found that beyond the storage capacity, initial states highly correlated with one individual pattern evolve to the corresponding class, while even the class categorization is lost at a higher critical loading [13]. In [14], it was proposed that such a scheme could function as a model for *prosopagnosia*, an impairment in visual recognition in which the patient can correctly recognize the category of faces but is unable to recognize individual faces.

However, can such a simple scheme be relevant to describe semantic memory, too? A tree-like structure, while possibly suited to capture a specific cognitive impairment, does not account for the complex relations of semantic memory. When dealing with the meaning of a concept, one typically accesses not only its identity and class membership, but also the stronger or weaker relations to other concepts, which span many dimensions and are not only contingent on common human experience but also on personal experience [15]. As such, the complexity of semantic relations [16] can be argued to require a more sophisticated description than the one provided by an approximate tree-like model.

Valuable attempts to go beyond both uncorrelated memories and simple branching trees, for example within the parallel-distributed processing (PDP) framework [17], have remained largely data driven, focused on computer simulations that could qualitatively reproduce results in agreement with patterns of deficit seen in the neuropsychological literature [18, 19, 20]. No mathematical framework, however, has been proposed for theoretical questions of a quantitative scope. Such a theoretical perspective is necessary if one wants to approach the question of semantic memory in a more principled way. For example, what is the reliability and generalizability of such results from the small networks used in simulations to large-scale cortical networks such as those of the human brain?

### 1.2 Connectivity

The analytical tools allowing for a complete analysis have been applied to fully connected or else very sparsely connected networks, in which the average connectivity between the units vanishes. These models have been thoroughly analyzed and scaling relations have been found for the storage capacity as a function of the mean connectivity and the coding sparsity in the network. Remarkably, such scaling relation holds, when coding is sparse, for both limit cases of full connectivity and extremely sparse connectivity. Does it mean that it holds also for any connectivity in between, including realistic models of cortical connectivity?

From the point of view of plausibility, such studies of randomly wired networks fall short of describing some features of the anatomy of cortical connectivity. For example, it has been shown [21] that in layers II and III of mouse visual cortex the probability of connection falls from 50 – 80 percent for directly adjacent neurons to 0 – 15 percent at a distance of 500 micrometers. Building on such observations, the properties of an autoassociative network of threshold-linear units whose synaptic connectivity is spatially structured has also been investigated [22]. Other experiments however, have shown that at a larger scale, cortical connectivity is not randomly distributed, not even after allowing for a distance-dependent parameter. For example, it has been shown that in the prefrontal cortex of monkeys, patches of a hundred microns make connections to and from other discrete patches of cortex of the same size [23]. A patch is connected to about 15 – 20 other patches in its proximity via grey matter connections, and to at least 15 – 20 more distant patches connected via white matter connections.

Braitenberg and Schuz have elegantly synthesized this dual local and global characteristic of the cortex in terms of the A and B systems (referring to apical and basal dendrites [24]). They suggest regarding the whole cortex as a memory machine, in which the B-systems encode a set of memories as local attractors and the A-system encodes global attractors, by virtue of long-range connections, Fig. 3(a). Variant models of associative memory networks that implement this separation of scale between dense local connectivity and sparse long-range connectivity have been studied [25, 26, 27, 28, 29]. This study is in line with such an approach, in that it aims at describing each patch of, say, the human cortex, a functional voxel of a few *mm*^3^, comprising some 10^5^ neurons, as one local network interacting through the B system, whose activity is coarsely subsumed into a Potts unit. A Potts unit has multiple activity states, akin to a *capsule* of the kind recently introduced in deep learning networks [30]. The Potts network, aimed at describing the cortex, or a large part of it, is comprised of *N* such units, constituting the A-system, Fig. 3(b). We refer to [31] for a detailed analysis of the approximate thermodynamic and dynamic equivalence of the full multi-modular model and the Potts network. We do not dwell on the correspondence here, but use it to discuss correlations in the Potts framework.

**Figure 3:**
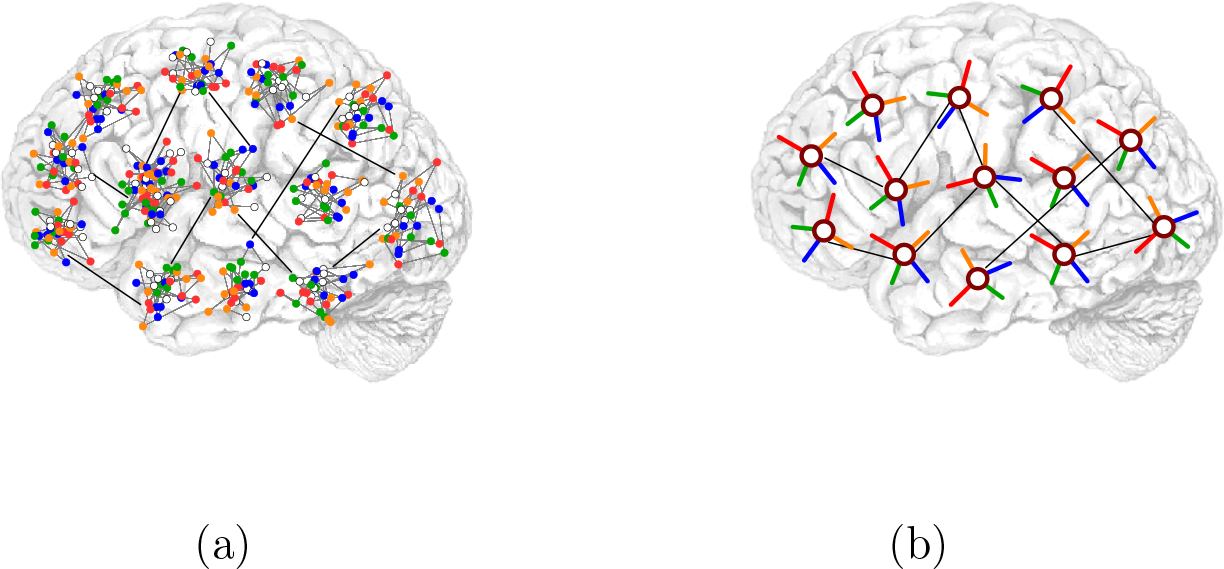
(**a**) The Potts network, here intended as a model of semantic memory, is a coarse description of the cortex in terms of local patches of dense connectivity, which store activity patterns corresponding to local attractors (**a**). Each patch is a small local network characterized by high connectivity; *diluted* connections are instead present between units of different patches. The configuration of the individual patch is assumed to converge to a local attractor, synthetically captured by a Potts state. Each Potts unit, depicted in (**b**) can be in any of S states, where green, orange, blue and red represent the active states (*S* = 4). The white circle at the center corresponds to the quiescent state, aimed at capturing a situation of no retrieval of the underlying local network.

## 2 Results

### 2.1 The Potts network

The Potts neural network is a generalization of Hopfield’s binary autoassociative network [32]. A Potts unit can be either in the *quiescent* state or in one of the S equivalent *active* states. By convention, we label these states with numbers from 0 to S, where k = 0 indicates the quiescent state and *k* = 1…S the active ones, representing the possible local attractors (see Fig. (3)). Due to stochastic fluctuations, a unit can be, with a non-vanishing probability, in several of the *S* + 1 states, so that the *activity* of unit *i* in state *k* is denoted by 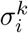, a variable in the interval [0,1]. By network state, or *configuration,* we refer instead to the collection of local states assigned to all units, {*ξ*_*i*_;}, each of which is an integer from 0 to *S*, and where *i* ∈ {1,… *N*}, *N* being the number of units in the network.

*Couplings J*_*ij*_ between states of distinct units represent the strength with which connected units influence each other. In the case of the Hopfield network, the couplings *J*_*ij*_ are just scalars. In the Potts network, the matrices 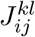 express the strength of the coupling between unit *i* in state *k* and unit *j* in state *l*.

Of crucial importance in the definition of the network model is the *learning rule,* which prescribes how the couplings depend on a given training data set. In the present model, the training data set consists of a number *p* of network configurations 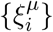 that we call *patterns*. Throughout this paper we consider patterns that are sparse: every unit in each pattern is taken to be in the quiescent state with probability 1 − *a*, with the remaining probability *a* shared uniformly by the *S* active states: 
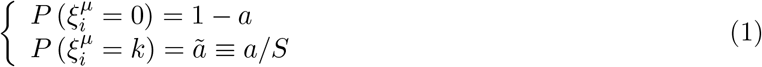

The way the patterns 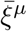 are generated, i.e. their probability distribution, has effects on the “retrieval properties” of the network, i.e. the ability to retrieve with good accuracy one of the training patterns, when it is partially cued. A quantitative measure of this ability of the network is the *storage capacity*, the number of patterns the network is able to store and retrieve, relative to the number of synaptic connections per unit.

The learning rule according to which the patterns are used to build the synaptic connections between units is a Potts-adapted version of Hebbian learning 
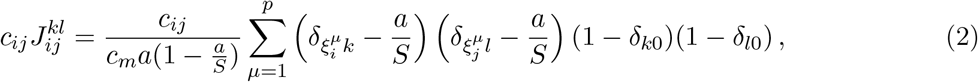
where the factor *c*_*ij*_ denotes the (*i*, *j*)-th entry of the adjacency matrix of the connectivity (graph), equal to 1 if an edge exists from *j* to *i* and 0 otherwise. The constant *c*_*m*_ is the average total degree of this graph, i.e. the average number of connections at a given node, so that ⟨*c*_*ij*_⟩ = *c*_*m*_/*N*. The Kronecker *δ*-function is 1 when the two indices are equal and 0 if they are different. The subtraction of the mean activity by state, *a/S*, ensures a higher storage capacity, as initially shown for the Hopfield network in [33] and for the Potts neural network in [34].

The fully connected network, in which *c*_*ij*_ = 1 for all pairs (*i*, *j*) allows for a full-fledged analytic approach, by means of techniques borrowed from spin glass physics [35]. It has been shown, as reviewed in [36], that such connectivity ensures that each of these configurations, if they are not too many, becomes a stable state, or an attractor of the energy function 
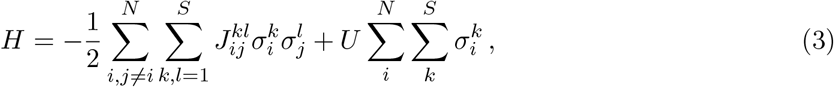
where the activation function is given by the Boltzmann distribution with inverse temperature *β* 
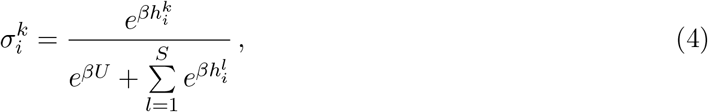
where *U* is a threshold for activation. The *field* received by unit *i* in state *k* is determined by the activity of all the Potts units and writes 
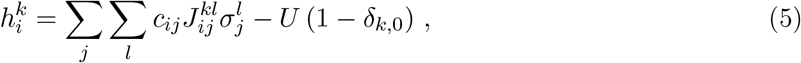
where the coupling strength between two states of two different units 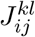 is given by Eq. (2). From Eq. (4), it follows that 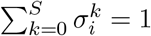 at all times.

A more biologically plausible case is that of *diluted* networks, where the number of connections per unit *c*_*m*_ is less than *N*. In this paper we consider *random dilution* (RD), in which 
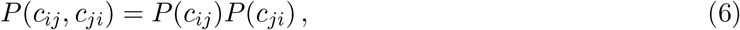
with 
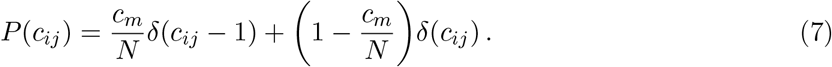

### 2.2 Generating correlated representations

The initial studies of the capacity of the Potts network [34, 31] featured patterns that were uncorrelated. *Uncorrelated patterns* are generated by assigning Potts states to different units in different patterns independently. This means that the *p* patterns 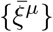 are generated according to a probability distribution which is factorized into *p* identical ones 
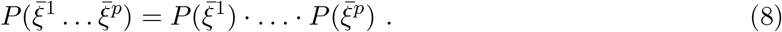

In turn, units in each pattern are also independent and identically distributed 
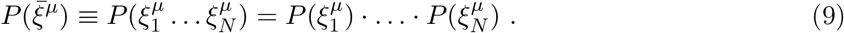

In this simple uncorrelated scheme, it is possible to compute the correlation between any two patterns *μ* = *v* as measured by the fraction of co-active units that are in the same state in both patterns 
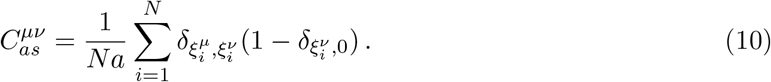

The distribution of this correlation measure is straightforward and given by a binomial 
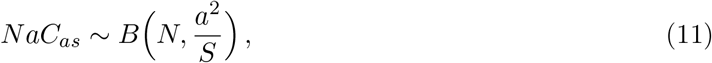
where 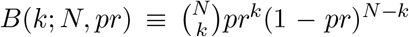; that is, ⟨*C*_*as*_⟩ = *ã*. Once patterns become correlated, however, there is no straightforward way to compute this measure, and we resort to simulations, as reported in Sect. 2.2.5.

#### 2.2.1 Single parents and ultrametrically correlated children

The interest in ultrametrically organized patterns was largely due to the discovery of an ultrametric hierarchy of the free energy minima in the formal solution of the Sherrington-Kirkpatrick model of a spin glass [35]. In particular, the Hopfield model of neural networks was extended to allow for the storage and retrieval of hierarchically correlated patterns [12]. In this study [12], a set of random patterns, which we can call “parents”, are characterized by independent units, active with probability *a* 
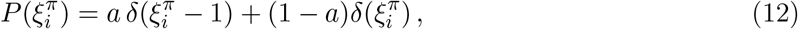
where 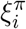 denotes the activity of unit *i* of parent *π* and 0 < *a* < 1 is the sparsity of the parents. In the next step, “child” patterns are drawn from the following distribution 
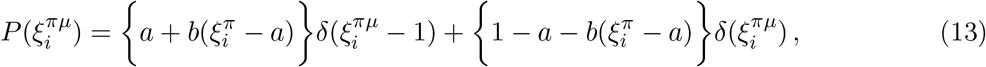
where 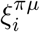 denotes the activity of unit *i* of child *μ* branching from parent *π*. 0 < *b* < 1 parametrizes to what degree children are biased toward their (single) parent. For *b* = 0, child patterns become uncorrelated with no dependence on the parent, while for *b* = 1 the child patterns become identical to their single parent. Given the distributions above, we can compute the average activity of parents and child patterns (since the state of each unit *i* is drawn identically from the same distribution, in the following we can drop this index) 
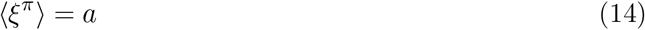

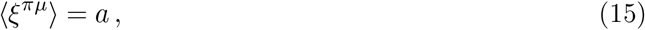
as well as child-parent correlations
 
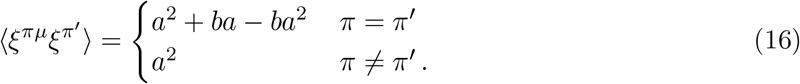

As expected, if *b* > 0, children of the same branch have higher similarity to their own parent (*π* = *π*′), than to a parent of another branch (*π* = *π*′). We can also compute the correlation between two children of the same parent (*π* = *π*′) and that of two children belonging to distinct parents (*π* ≠ *π*′) 
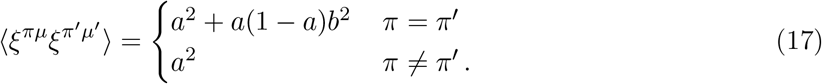

It trivially follows that 
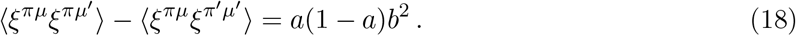

This is one of the characteristics of this algorithm: it is possible to define a distance *d* such that three patterns (*x*, *y*, *z* = *ξ^πμ^*, *ξ^πμ′^*, *ξ^πμξ^*) *at the same level of the hierarchy* can be seen to satisfy the *strong triangle inequality*: *d*(*x*,*z*) ≤ max(*d*(*x*, *y*), *d*(*y*, *z*)) and permutations of *x*,*y*,*z*. As illustrated in Fig. 4(a), triplets of patterns can only be in one of the two triangle relations: equilateral and isosceles with two long edges, in other words, an ultrametric space has no node intermediate between any two nodes (Fig. 4(b)).

**Figure 4:**
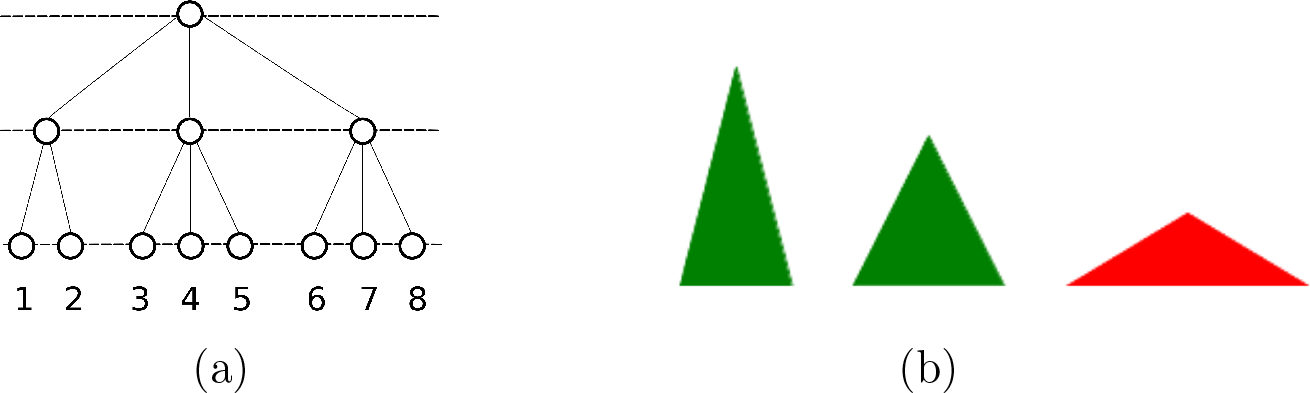
(**a**) A tree, adapted from [35]. 1, 2, 3, 4, 5, and 6 are at the same level of the hierarchy. If we consider the nodes 1, 3 and 6, they are each at a distance of 2 of each other, the distance being defined as the distance to the nearest common branching point. If we consider nodes 3, 4 and 5, then they are each at a distance of 1 of each other, such that we get again an equilateral triangle. If we consider 1, 2 and 3, then *d*_12_ = 1 while *d*_13_ = *d*_23_ = 2, such that we get an isosceles triangle with two long edges. One alternative, an isosceles triangle with two short edges, is impossible to realize: there are *no intermediate points* between 1 and 3 or 2 and 3, as indicated in red in (**a**).

From the point of view of semantics, this is an implausible situation. If one considers superordinate categories as the single archetypal parent from which all concepts descend, it becomes clear that such an ultrametric structure is unsuitable in describing all the semantic relations in which the ultrametric inequality is not satisfied: for example when a concept finds itself “in between” two other concepts. On the other hand, the very meaning of a concept can be thought of as the set of features that are associated to it. It may then be more sensible to consider the features characterizing a concept as its building blocks, hence its parents. In the following, we describe an algorithm, first sketched in [37], in which each child pattern (concept) is generated from multiple parents (features), a random subset of the total group of parents relevant to it.

#### 2.2.2 Multiple parents and non-trivially organized children

How can we incorporate a plausible featural description into our model of semantic memory? One may consider features as the parents from which child concepts are derived. We can then map quantities such as the number of features, their sharedness, and their dominance to appropriate parameters in our model.

Our simple version of the multi-parent pattern generation algorithm works in three stages. In the first stage, a set of Π random patterns are generated to act as parents. In the second stage, each of the Π parents are assigned to *p*_*par*_ randomly chosen children. Then, each “child” pattern is generated: each pattern, receiving the influence (or input) of its parents, aligns itself, unit by unit, in the direction of the largest input. In the third and final step, the fraction a of the units with the largest inputs is set as active in each child pattern. A schematic representation can be seen in Fig. 5(b) and put in contrast with a schematic representation of the single parent algorithm in Fig. 5(a).

**Figure 5:**
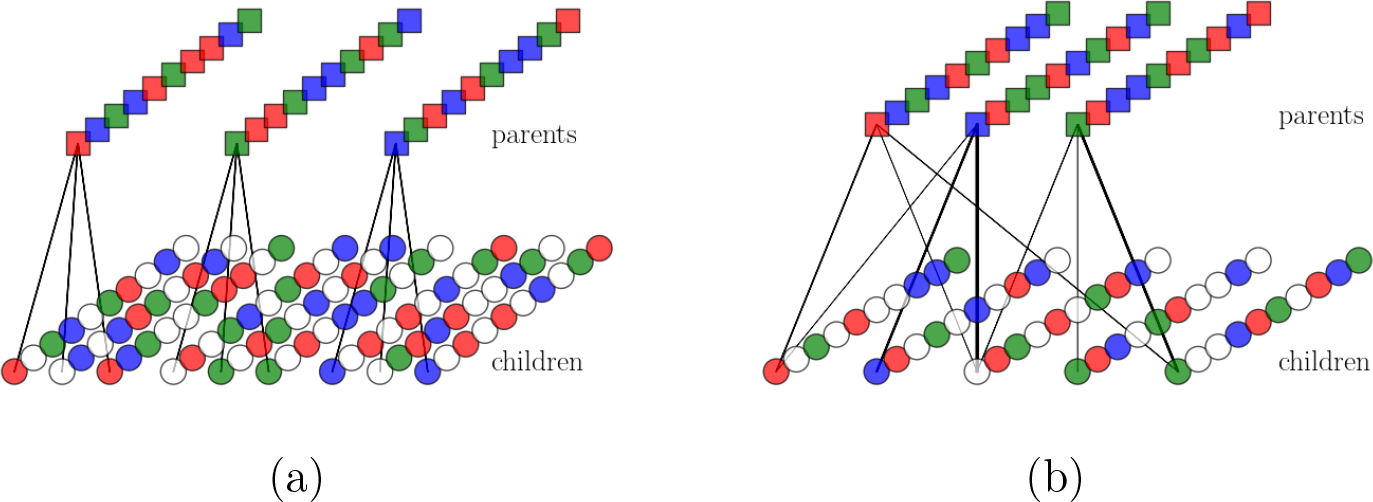
(**a**) The workings of a hierarchical algorithm with 3 parents and 3 child patterns per parent. The different lines of squares/circles correspond to the different units in parents/children. Colors correspond to active Potts states while white denotes the quiescent states. *S* = 3. (**b**) The workings of the multi-parent algorithm with *Π* = 3 parents and *p*_*par*_ = 3 child patterns per parent and 5 total child patterns. Black arrows and their thickness denote strength of input. The main difference with the hierarchical algorithm is that each child pattern can receive input from multiple parents. If each parent is to represent a feature and each child a concept, the algorithm entails the generation of a concept from multiple features.

#### 2.2.3 The algorithm operating on simple binary units

Each parent is assigned *p*_*par*_ children out of a total of *p*. The probability distribution that a given child has *n*_*p*_ parents, out of a total pool of Π is given by a binomial, with the *prolificity f* = *p*_*par*_/*p* 
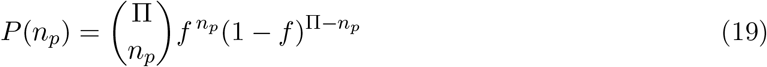

The algorithm draws, for the input 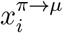 from unit *i* of parent *π* to unit *i* of pattern *μ*, a uniformly distributed random number in the interval (0,1] with probability *a*_*p*_ and zero with probability 1 – *a_p_* such that we can write 
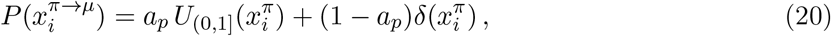
where *a*_*p*_, which we can call the *extent* of the input from one parent, is analogous to the a parameter in Eq. (12); indeed, if *a*_*p*_ ~ 0, a child pattern is very unlikely to receive, on a particular unit, the contribution from one of its parents. On the other hand, if *a*_*p*_ ~ 1 then all parents influencing a child contribute to its input, whichever the unit. *U*_(0,1]_ denotes the uniform distribution, such that the input from parents is graded, contrary to the previous section.

Here, we have made the choice of non-sparse parents, but sparse input from parents, aimed at decorrelating units, while conserving correlations between patterns. This choice will prove crucial in Sect. 2.3.1, where statistical independence between units will lead to a vanishing mean noise, using only a simple covariance rule. For *S* = 1, this means that the patterns generated by the algorithm are uncorrelated, but the importance of having non-sparse parents with sparse input from them becomes important when dealing with more than one Potts state. Nevertheless, in this section, we consider *S* = 1, before treating genuine Potts units.

The main difference with respect to the single-parent algorithm is that now, one must compute the total field 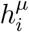 that a unit *i* of pattern *μ* receives from all parents 
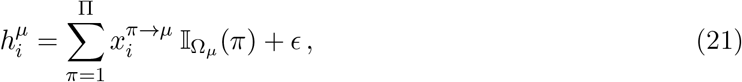
where Ω_*μ*_ is the set of all parents acting on pattern *μ* and where we have 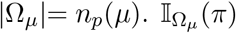 is the indicator function that is 1 if parent *π* is assigned to pattern *μ* and 0 otherwise. *ϵ* is a small random input (*ϵ* ≪ 1) allowing for some input, even when *a*_*p*_ ≪ 1. The fields of all units of all patterns have the same distribution. In App. A, the full derivation of the probability distribution for the field 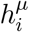 is reported. Such a distribution has a non-trivial expression and, to our knowledge, it can only be evaluated numerically. However, a simple analytic expression can be given for the moments of the distribution of 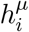 
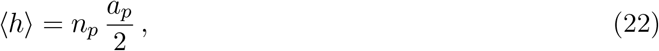

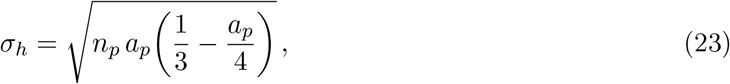
as shown in Fig. 6(b). In Fig. 6(a), we see that these analytical results match tightly those from implementation of the algorithm.

**Figure 6:**
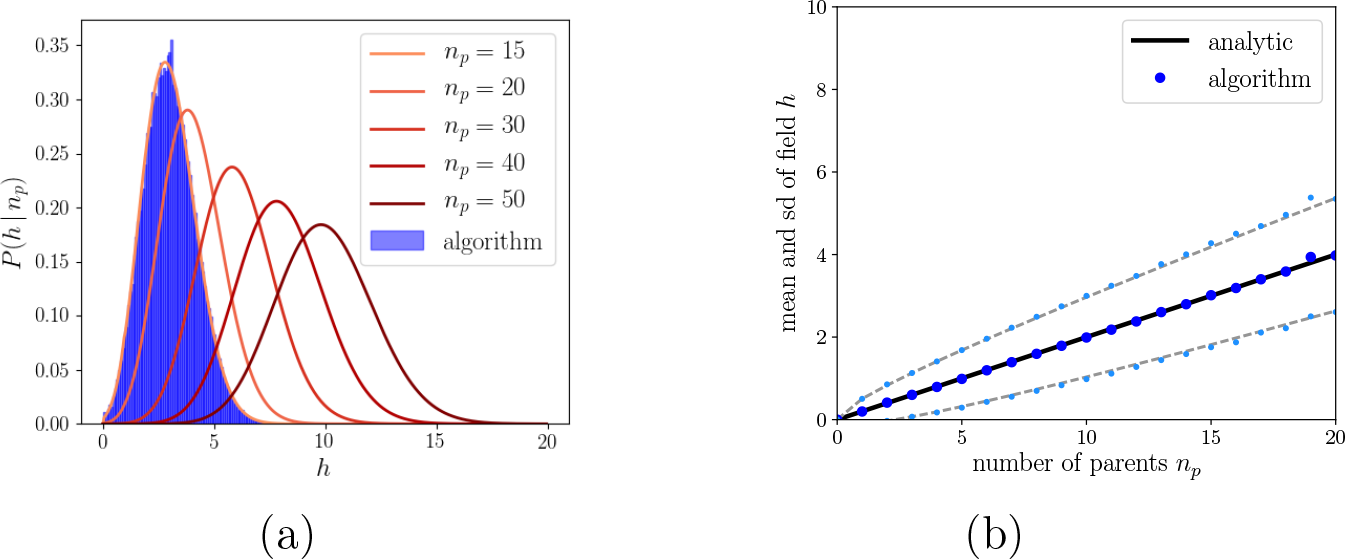
(**a**) Solid lines correspond to the analytical distributions of the field, Eq. (A8), in blue is the distribution of the fields produced by a simulation of the algorithm for *n*_*p*_ = 15. The parameters are *N* = 2000, *S* = 1, *a*_*p*_ = 0.4, *n*_*p*_ = 15 … 50 and Π = 100. (**a**) The mean and standard deviation of the field as a function of the number of parents.

As a last step, within a given pattern, a fraction *a* of the units having fields above a threshold *h*_*m*_ are set to become active. The threshold *h*_*m*_ is then implicitly given in terms of the cumulative distribution function 
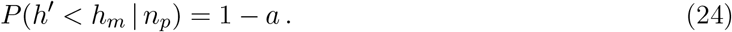

For any given child pattern *μ* with number of parents *n*_*p*_, we can now define the probability that it will be activated, given the field that it receives 
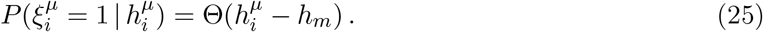

#### 2.2.4 The algorithm operating on genuine Potts units

With genuine Potts states, the main difference with respect to the previous case is that the input from a parent *π* to the field of its child patterns can be, on a given unit, to any one of *S* states, with equal probability. This means that only a subset Ω_*i*, *k*_ of the total parents will contribute to state *k* of unit *i*. We denote the number of parents in the subset as 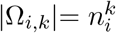. The joint distribution of number of parents by state is 
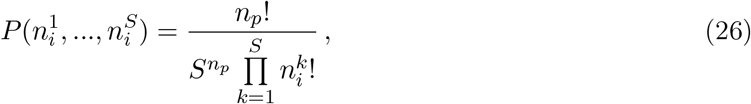
such that the constraint 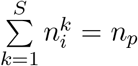 is satisfied. We can then write the field of unit *i* in state *k* of pattern *μ* 
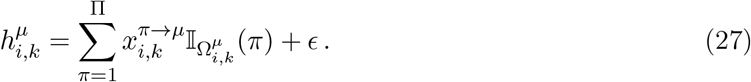

Then, the algorithm is such that it selects, unit by unit, the state receiving the maximal input. Following some calculations shown in App. B, we can compute the distribution of the fields for those states having received maximal input *H* (Fig. 7(a)). We can then compute, exactly as before, the threshold above which the unit becomes activated 
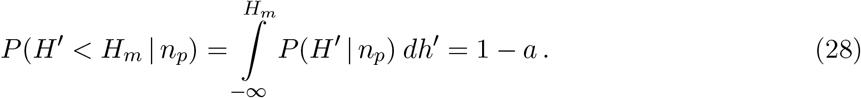

**Figure 7:**
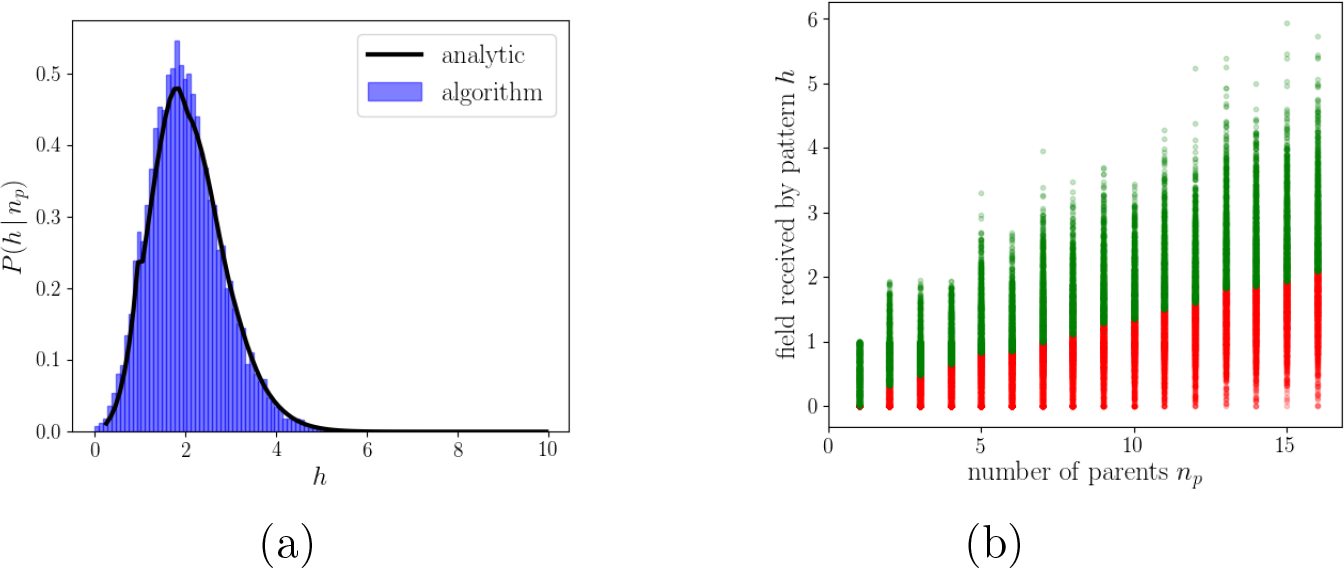
(**a**) Distribution of the maximal fields for *S* = 2 and *n*_*p*_ = 30. In blue is the distribution of the fields produced by the algorithm and the black line is Eq. (B7). (**b**) The *x*-axis orders patterns with different number of parents and the *y*-axis the fields of the units in that pattern. Red points correspond to units that are set to quiescent and green to those that are activated. The boundary between the green and the red corresponds to *h*_*m*_, the minimum field required for a unit to be set to active. Parameters are *N* = 2000, *S* =2, *a*_*p*_ = 0.4 and Π = 100.

Having obtained the minimal field *H*_*m*_ required to activate a unit (Fig. 7(b)), we now need only the distribution of the field given the number of parents in that state *P*(*h*^*k*^~*n*^*k*^), which is none other than Eq. (A8) (replacing *n*_*p*_ with *n*^*k*^). We finally get to the distribution of activity across units and states, given the field received 
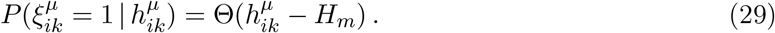

Given the algorithm just described, the main mechanism determining the state of a unit in a given pattern is how many of the parents affecting a child are in the same state. If parents are all aligned, this makes the unit receive a higher field in a single state, making it more likely to become activated. On the other hand, lower alignment between parents results in the field received by a child unit to be spread among the different states, and make it less probable for the child unit to find itself among those with maximal fields, as given by Eq. (B7).

#### 2.2.5 Resulting patterns and their correlations

In the previous section, we have described the mechanism through which individual child patterns are generated. At this level, in order to determine whether or not a unit of a pattern is active, the only relevant parameter is the number of parents, as well as their degree of alignment in Potts space. From the point of view of an individual child pattern then, all parents are equivalent and can be considered as identical and independently distributed, a property exploited above. In this section, we turn to the correlations between patterns. Are they dominated by the number of parents that a pair of child patterns have in common? Is this a plausible model for semantic memory?

Patterns generated by the algorithm sample different active states uniformly, such that Eq. (1) still holds, though the joint distribution 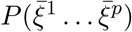 is not factorizable anymore, as it was in Eq. (8). In Sect. 2.2.2 we discussed how the activity of different units is still approximately uncorrelated. We can see this by computing, analogously to the correlation between patterns, Eq. (10), the correlation between units as the fraction of patterns in which two units are co-active and in the same state 
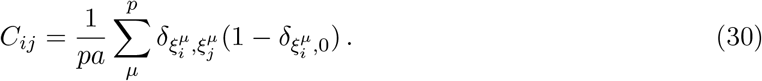

In Fig. 8 we can see the distributions of *C*_*μv*_ and *C*_*ij*_ for nine different combinations of the extent *a*_*p*_ and prolificity *f* parameters. The distributions are very sensitive to the specific values of the parameters. For low values of *a*_*p*_ and *f*, pairs of Potts units have uncorrelated activity when averaged across patterns, in the sense that the distribution *C*_*ij*_ has zero covariance. Pairs of patterns, instead, are correlated with a distribution *C*_*μv*_ of non-zero covariance, that is positively skewed. Low values of *a*_*p*_ and high values of *f* result in the distribution of the pattern correlations becoming more and more normal, while high values of *a*_*p*_ and low values of *f* result in a normally distributed correlation between units and a highly skewed multi-modal distribution between patterns.

**Figure 8:**
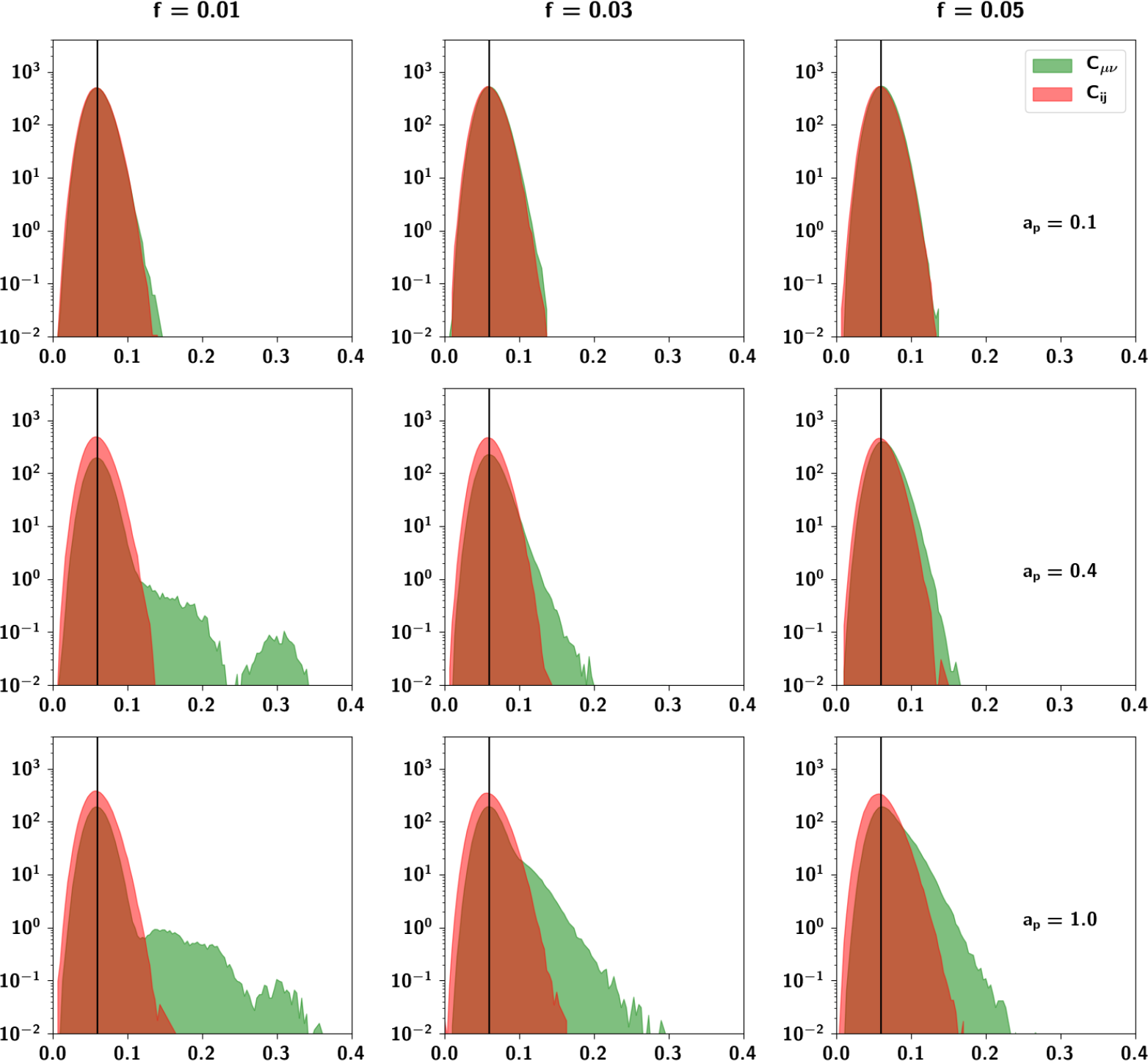
Probability density function of correlations between units (in red) and between patterns (in green) for three different values of both *a*_*p*_ and *f*, the latter yielding in this case an average of 1.5, 4.5 and 7.5 parents per pattern. The black vertical line corresponds to the average correlation with uncorrelated patterns distributed independently according to Eq. (1). The parameters are *S* = 5, *a* = 0.3, and Π = 150. The algorithm produces correlations between patterns with high variability relative to the correlation between units, in line with ideas about semantic memory. Note that the algorithm is sensitive to the parameters and their values strongly affect the correlation between patterns.

To assess these observations more systematically, in Fig. 9(a) we can see boxplots of the *C*_*μv*_ distributions for different values of *a*_*p*_ keeping *f* = 0.05 fixed. While the mean correlation is unaffected by increasing *a*_*p*_, the standard deviation and the skewness increase. In 9(b), conversely, we can see boxplots of *C*_*μv*_ distributions for different values of *f* keeping *a*_*p*_ = 0.4 fixed. It can be seen that increasing *f* increases the mean correlation between patterns. The effects observed can be understood intuitively because of the different roles that these parameters play in the algorithm. The extent *a*_*p*_ is the parameter that increases the probability that a child unit receives input from a parent, increasing the overall similarity of a child to its parents. This means that those children that have a larger number of shared parents will be more similar and more strongly correlated, giving rise to the larger values in the distribution. The prolificity *f*, on the other hand, is the ratio of the pool of children affected by one parent to the total number of children. Increasing this ratio leads to an increase in ⟨*n*_*p*_⟩, the mean number of parents, such that children tend to share more parents. It can be seen in Fig. 10(b), in which pairs of patterns are decomposed into different distributions sharing an increasing number of parents (0 – 5 shared parents), that for a pair of patterns, a higher number of shared parents leads to a higher mean correlation. The number of such pairs is markedly fewer, as can be seen in the left axis of Fig. 10(a) (plotted in a logarithmic scale), but if f is high enough, this effect is enough to increase the overall mean correlation between all patterns. The two parameters *a*_*p*_ and *f* therefore play different roles in generating the correlations.

**Figure 9:**
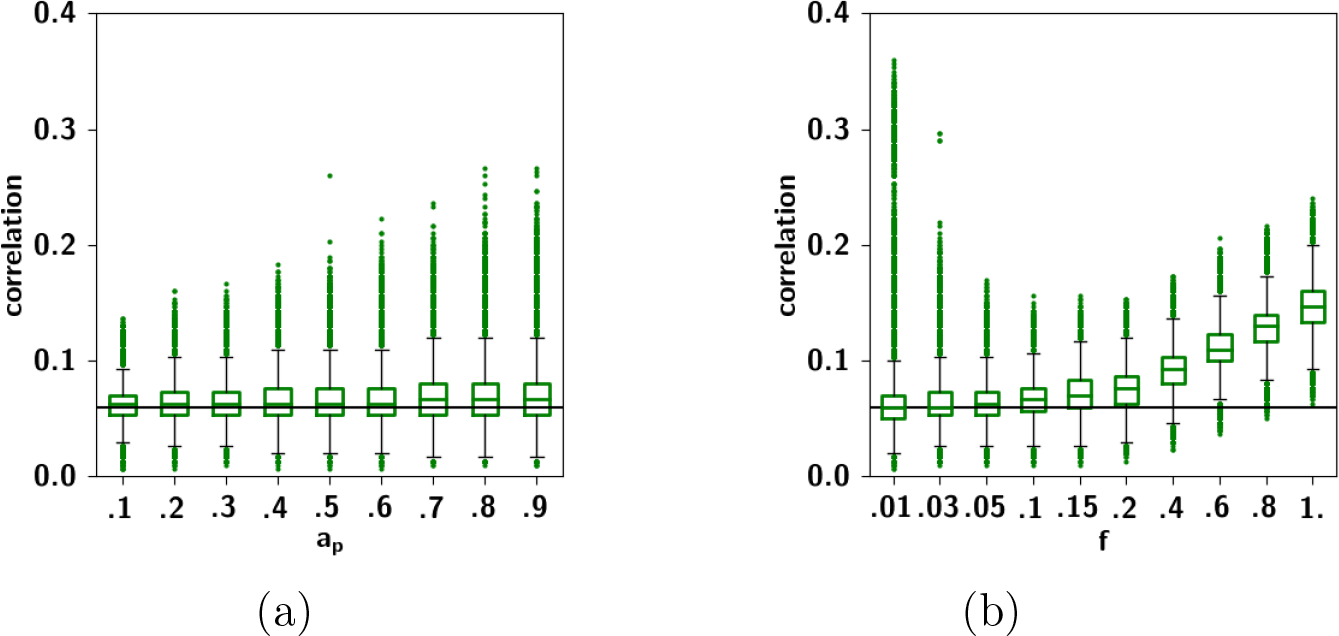
(**a**) Boxplots of *C*^*μv*^ for different values of *a*_*p*_, with *f* = 0.05 fixed. (**b**) Boxplots of for different *C*^*μv*^ values of *f* with *a*_*p*_ = 0.4 fixed. The parameters *a*_*p*_ and *f* play different roles in generating the correlations. Increasing the extent *a*_*p*_ of the input they receive from each parent increases the overall similarity of those children having shared parents, as evidenced by the increasing skewness of the distributions. In contrast, increasing the prolificity *f*, leads to an increase in the mean number of shared parents, such that all children are more correlated, as shown by the shift in the overall distribution. The black horizontal line corresponds to the average correlation with uncorrelated patterns distributed according to Eq. (1). Other parameters are *a* = 0.3, *S* = 5 and Π = 150.

**Figure 10:**
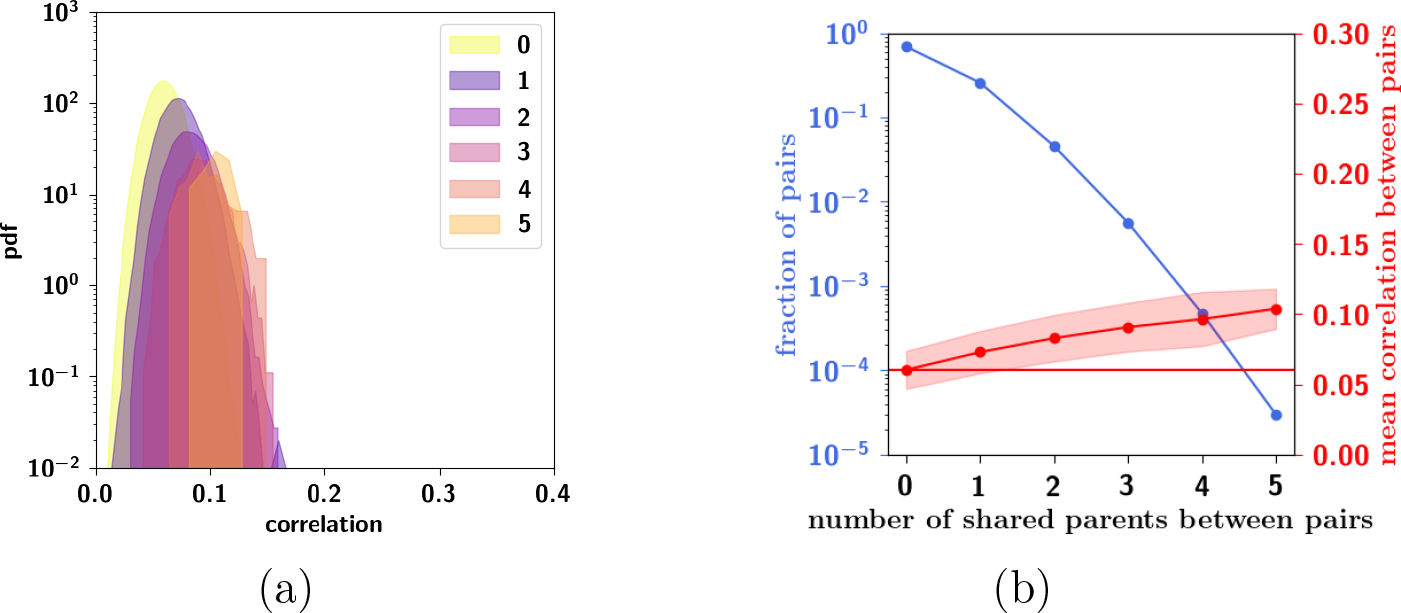
(**a**) Another visualization of the correlation distribution of Fig. 8, with *f* = 0.05, *a*_*p*_ = 0.4 and Π = 150, decomposed into the distribution for each number of shared parents. (**b**) Fraction of pairs of patterns (left *y*-axis, note the logarithmic scale) and mean correlation between those pairs (right *y*-axis, linear scale) as a function of number of shared parents. The red horizontal line corresponds to the average correlation with uncorrelated patterns distributed according to Eq. (1). Pairs of patterns having more shared parents are markedly fewer, although on average more correlated, so they do not affect much the overall mean correlation.

#### 2.2.6 The ultrametric limit

It is interesting to note a limit case of the algorithm. For low prolificity, if e.g. ⟨*n*_*p*_)_μ_ = Π *f* ~ 1 as in Fig. 8 (left column, i.e., *f* = 0.01, Π = 150), on average most children will have a single parent, which effectively produces ultrametric patterns. Indeed, for these parameters, since the number of total parents Π = 150 is smaller than the total number of children generated, *p* = 1000, several children share a given single parent. The mean value of their correlation with all other children, however, at *a/S*, is the same as the mean correlation between uncorrelated patterns. Note that the distribution is multimodal. The values forming the second mode of the distribution express the correlation between children belonging to the same (single) parent.

#### 2.2.7 The random limit

Another limit is the random or limited-parent-influence limit, in which *a*_*p*_ ≪ 1 (effectively, the top row in Fig. 8). In this case, most units will not receive input from their respective parents, regardless of how many they are, and the unit will align itself in the direction of a random Potts state given by the input *ϵ*. In this way, it is possible to parametrically generate patterns ranging from independent (*a*_*p*_ ≪ 1) to ultrametric (*a*_*p*_ = 1, *f* Π = 1), from the top row to the left column of Fig. 8, but also to enter the area of complexity to the bottom right, where correlations might begin to resemble plausible semantic relations.

#### 2.2.8 Semantic dominance

Returning to the correlation observed among the nouns we considered in Sect. 1.1 as our toy example, how important is, in the set of words, each individual feature? We can quantify it through a simple measure of semantic “dominance”, by simply summing the feature weights of all nouns 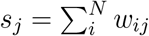

In Fig. 11, we report the summed weights of the *M* = 50 features across all the nouns considered, sorted and plotted on a semi-logarithmic scale. Even given the very small dataset used, it can be approximated to a good extent by an exponential law. The suboptimal fit may conceivably be the result of limited and unbalanced sampling. Indeed, the words, the nouns or the verbs were not chosen with comparable frequency. This measure is therefore only approximate, as an aggregate measure of dominance. Our measure is related to the measure called “semantic relevance” used by Sartori and colleagues [38] as well as to the “semantic differential” used by Osgood [39]. The difference with the latter measure, however, is that ours is cumulative across all of the nouns and derived from co-occurrence statistics in a corpus, while the semantic differential refers to a scale in which individuals rate the connotative meaning of objects, events, and concepts.

**Figure 11:**
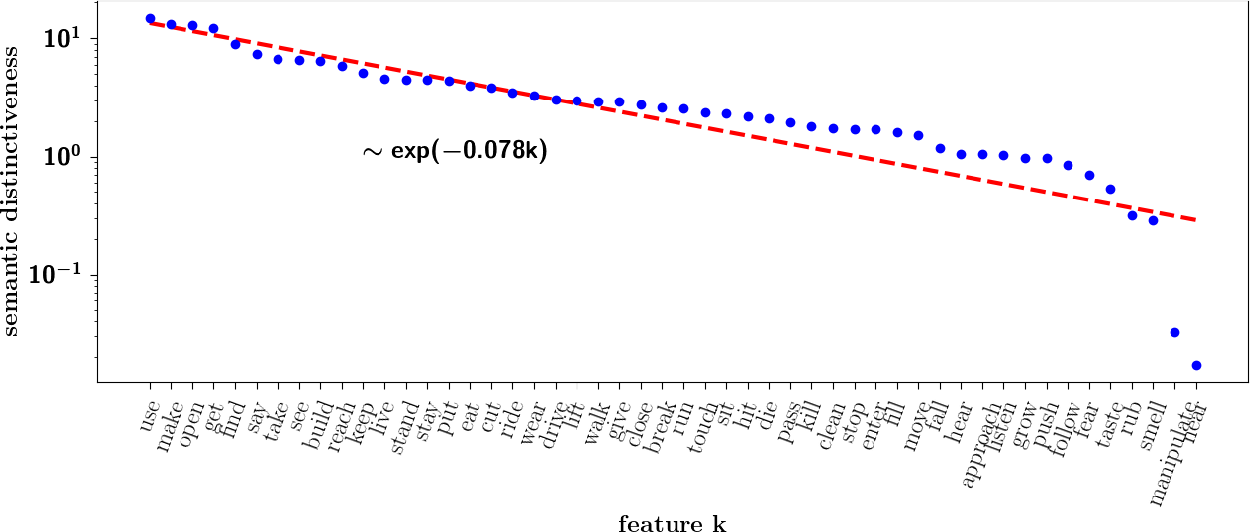
The *x*-axis lists all of the features used to compute the correlation between the nouns in the toy example of Sect. 1.1, sorted according to their summed weights across all nouns (reported on a semi-logarithmic *y*-axis). The exponent of the fit, *ζ* = 0.078, indicates that the semantics of this particular set of nouns is effectively dominated by a set of order 1/*ζ* ≃ 10 features.

To take into account this observation, we consider a more refined model in which the parents in our algorithm (the features), ranked from 1 to Π, have their input strengths damped exponentially with a dominance rate *ζ*, such that Eq. (27) is revised in the following way 
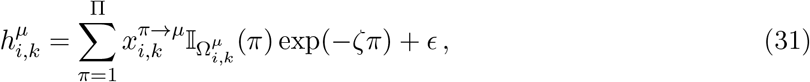
where 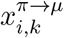 is the input from parent *π* to child pattern *μ*, 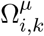 is the set of all parents acting on pattern *μ* and 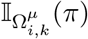 is the indicator function that is 1 if parent *π* is assigned to pattern *μ* and 0 otherwise. The limit *ζ* → 0 corresponds to the algorithm described in the previous sections, such that we recover Eq. (27). In this way, we introduce a parameter, *ζ*, which can be related to the slope seen in dominance distributions observed in real data, such as the one in Fig. 11.

In Fig. 12 we can see a schematic representation of this new algorithm. In contrast to the extent *a*_*p*_, the parameter *ζ*, though also affecting the strength of input, plays a different role, as it affects the global strength with which each parent affects its children, leading to variability of input across patterns. A high value of *ζ* contributes to highly unbalanced input from parents influencing a child pattern, such that units tend to align each with the most powerful parent, or the most dominant feature.

**Figure 12:**
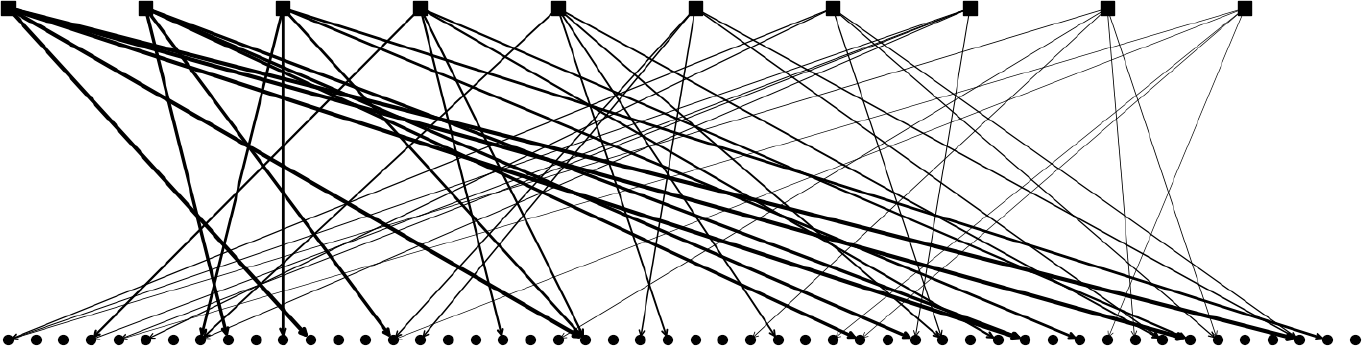
One sample representation of parent-child relations. The squares on the top row represent parents, while the circles at the bottom row represent children. Black lines represent input from the parents to the children. The strength with which each parent affects its children is proportional to exp(−*ζπ*), where π indexes the parents, as explained in the text. For illustration, there are Π = 10 parents, *p*_*par*_ = 5 children per parent and *p* = 50 total children.

How are the correlations affected by the dominance *ζ*? In Fig. 13 we report the distributions for three different values of the dominance *ζ* and prolificity *f*. While for low values of *ζ*, i.e. parents homogeneous in their strengths, the correlation between patterns is unaffected (see Fig. 8), increasing *ζ* we see the emergence of a tail of highly correlated patterns. For small *f*, this has the effect of smearing the bi-modal distribution, while for larger *f*, the already existing tail becomes fatter. This effect can also be seen more summarily in Fig. 14. Parents’ dominance reinforces children correlations such that in the regime of low dominance, we call the resulting patterns *weakly correlated,* while in the regime of high dominance, we call the resulting patterns *strongly correlated*.

**Figure 13:**
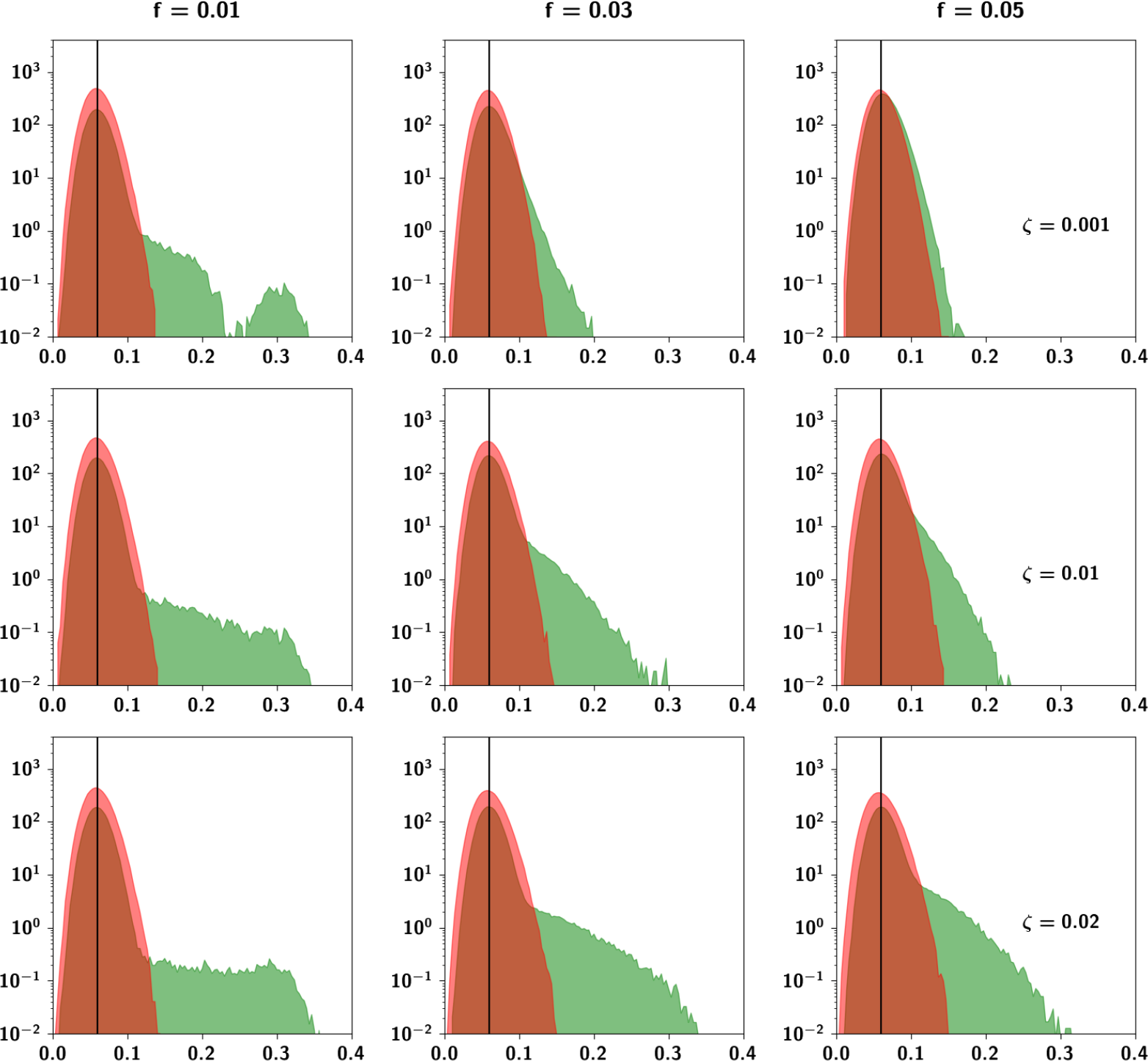
Probability density function of correlations between units (in red) and between patterns (in green) for three different values of the dominance rate *ζ* and prolificity *f*, keeping *a*_*p*_ =0.4 constant. For the low value of *ζ* = 0.001, this figure reproduces the middle panel of Fig. 8. For higher values of *ζ*, where the parents become highly heterogeneous, we see the emergence of large correlations.

**Figure 14:**
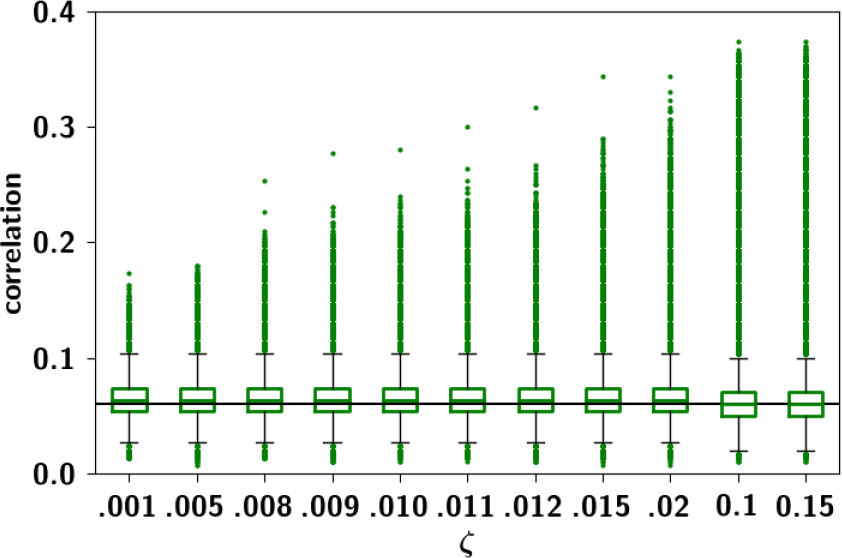
*C*^*μv*^ for different values of *ζ*, with *a*_*p*_ = 0.4 and *f* = 0.05 fixed. Other parameters are *a* = 0.3, *S* = 5 and Π = 150.

### 2.3 Storage capacity of the Potts network with correlated patterns

Having defined an algorithm which generates correlated patterns, we can turn to study the storage capacity of the Potts network and how it is affected by the correlations. We have carried out numerical simulations [34] with the learning rule in Eq. (2), and have observed that the storage capacity is diminished in the case of correlated patterns, a result that has been obtained analytically by others [40, 41], albeit for different sources of correlations.

#### 2.3.1 Self-consistent signal to noise analysis

In [31], we have discussed the application of the self-consistent signal to noise analysis (SCSNA) to the Potts network with uncorrelated patterns. In the following section we extend this analysis to the case of correlated patterns encoded by a Potts network with diluted random connectivity (Eq. (7)) and obtain estimates of the storage capacity accounting for correlations. In the expression of the field, Eq. (5) we identify two terms which are commonly referred to as signal and noise, with respectively non-vanishing and vanishing averages. While the signal, the contribution from the condensed pattern (that we label as *μ* = 1 in Eq. (2)), is what pushes the activity of the unit such that the network configuration converges to an attractor, the noise, or the crosstalk from all of the other patterns, is what deflects the network away from the cued memory pattern. Denoting 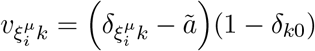 the noise term writes 
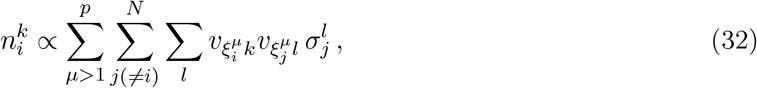
that is, the contribution to the weights 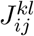 by all non-condensed patterns (*μ* > 1). By virtue of the
subtraction of the mean activity in each state *ã*, the noise has vanishing average: 
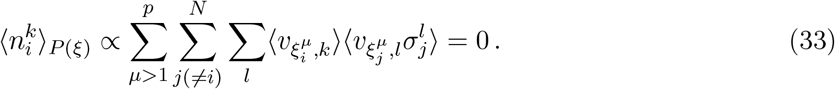

The variance of the noise can be approximately written in the following way: 
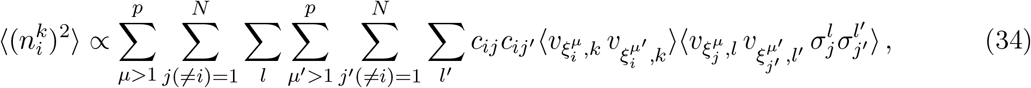
where statistical independence between units is implicitly used. While in the case of uncorrelated patterns, all terms but *μ* = *μ′*, *j* = *j′* and *l* = *l′* vanish, with correlated patterns this is not the case. Now, the additional terms *μ* ≠ *μ′*, *j* = *j′* and *l* = *l′* must be considered. Given the statistical
independence of units, however, all other terms are zero. Having identified the non-zero terms, we can proceed with the capacity analysis. We can express the field, Eq. (5) using the overlap parameter 
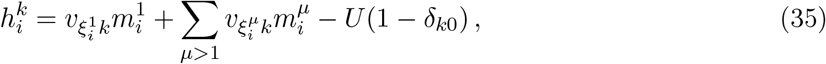
where we define the local overlap 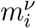 as 
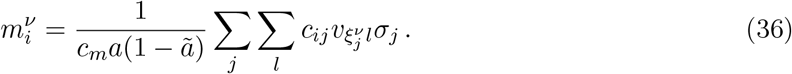

At the root of the SCSNA [42, 22, 43] is the assumption that the noise term itself can be expressed as the sum of two terms, one proportional to the activity of unit i and the other a Gaussian random variable, 
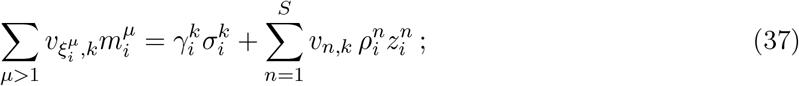
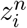 are standard Gaussian variables, and 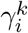 and 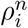 are positive constants to be determined self-consistently. The first term, proportional to 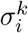, represents the noise resulting from the activity of unit *i* on itself, after having reverberated in the loops of the network; the second term contains the noise which propagates from units other than *i*. The activation function writes 
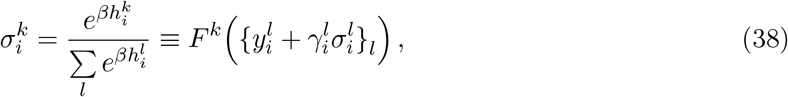
where 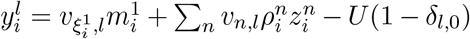. The activity 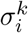 is then determined self-consistently as the solution of Eq. (38) 
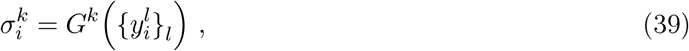
where *G*^*k*^ are functions solving Eq. (38) for 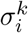. However, Eq. (38) cannot be solved explicitly. Instead we make the assumption that 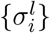 enters the fields 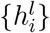 only through their mean value 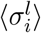, so that we write 
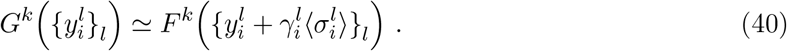

The coefficients in the SCSNA ansatz, Eq. (38), 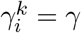 and 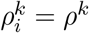 are found to be 
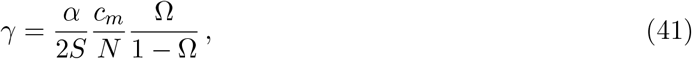

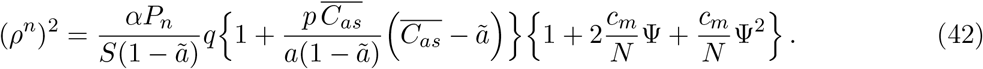
where *P*_*n*_ refers to the distribution of the patterns (Eq. 1, where *α* = *p/c*_*m*_ as before, and where *C*_*as*_, defined in Sect. 2.2.5, is the fraction of units that are in the same Potts state in two different patterns, normalized by *a*. Note the second term in the first curly brackets that scales with *p*^2^/*c*_*m*_ and is proportional to 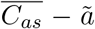 the covariance between patterns. This term originates from the additional non-zero terms in the sum in Eq. (34) due to correlations between patterns. When uncorrelated patterns are considered, such that 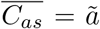, it becomes zero. In this calculation, we assume that correlations between the *v* operators of order higher than the second are negligible. As a consequence, the only quantities involved are their covariances. This approximation corresponds to the assumption that the *v* operators are normally distributed. Following the same procedure reported in [31], Ω, *q* and Ψ are found to be 
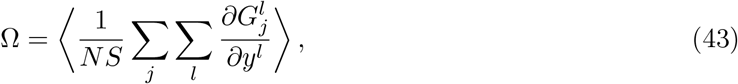

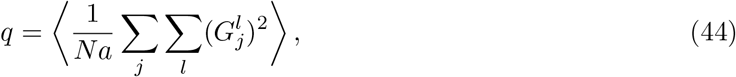

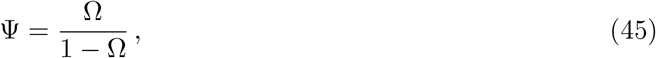
where ⟨·⟩ indicates the average over all patterns. The mean field received by a unit is then 
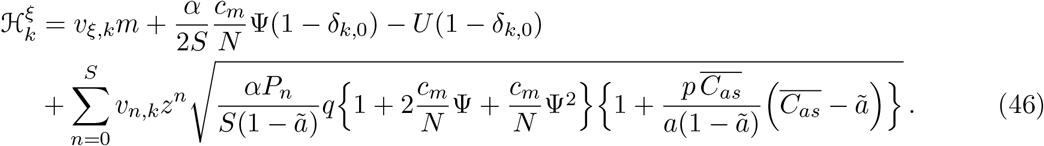

Taking the average over the non-condensed patterns (the average over the Gaussian noise *z*), followed by the average over the condensed pattern *μ* = 1 (denoted by ⟨·⟩_*ζ*_, in the limit *β* → ∞ we get the self-consistent equations satisfied by the order parameters 
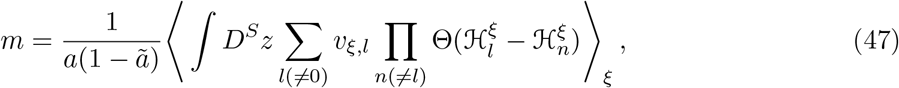

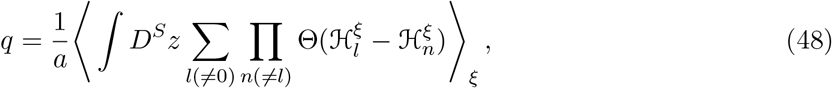

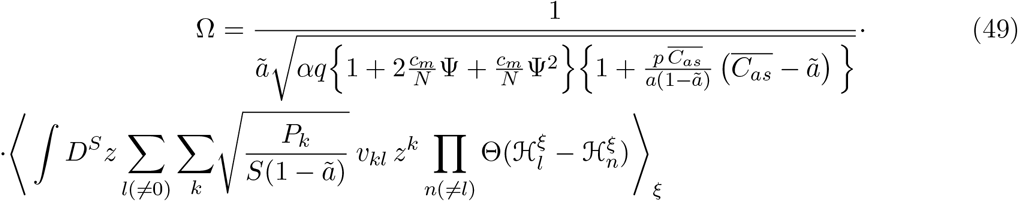

The averaging in Eqs. (47)-(49) can be performed analytically and we refer to [34] and [31] for their expressions. The storage capacity *α*_*c*_ is defined as the maximal *α* that solves the set of equations Eqs. (47)-(49) with finite overlap *m*.

#### 2.3.2 Numerical solutions of mean-field equations and simulations

In Fig. 15(a) we solve the set of self-consistent equations Eqs. 47-49 for different values of the sparsity and *c*_*m*_/*N* for the simpler case of uncorrelated patterns. In Fig. 15(b) we can see the agreement of the former solutions for *c*_*m*_/*N* = 0.1 with simulations. We can also see, in the same figure, the mean-field solutions as well as simulations for correlated patterns, with the values of 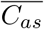 obtained from simulations of the algorithm. For lower values of the sparsity, the solution to the mean-field equations over-estimates the capacity compared to what we obtain through the simulations, possibly because the mean-field treatment does not account for the fluctuations in the correlations obtained through the algorithm, but only the increase in the excess mean correlation. For higher values of the sparsity, the agreement is better presumably because the correlations produced by the algorithm become dominated by the mean.

**Figure 15:**
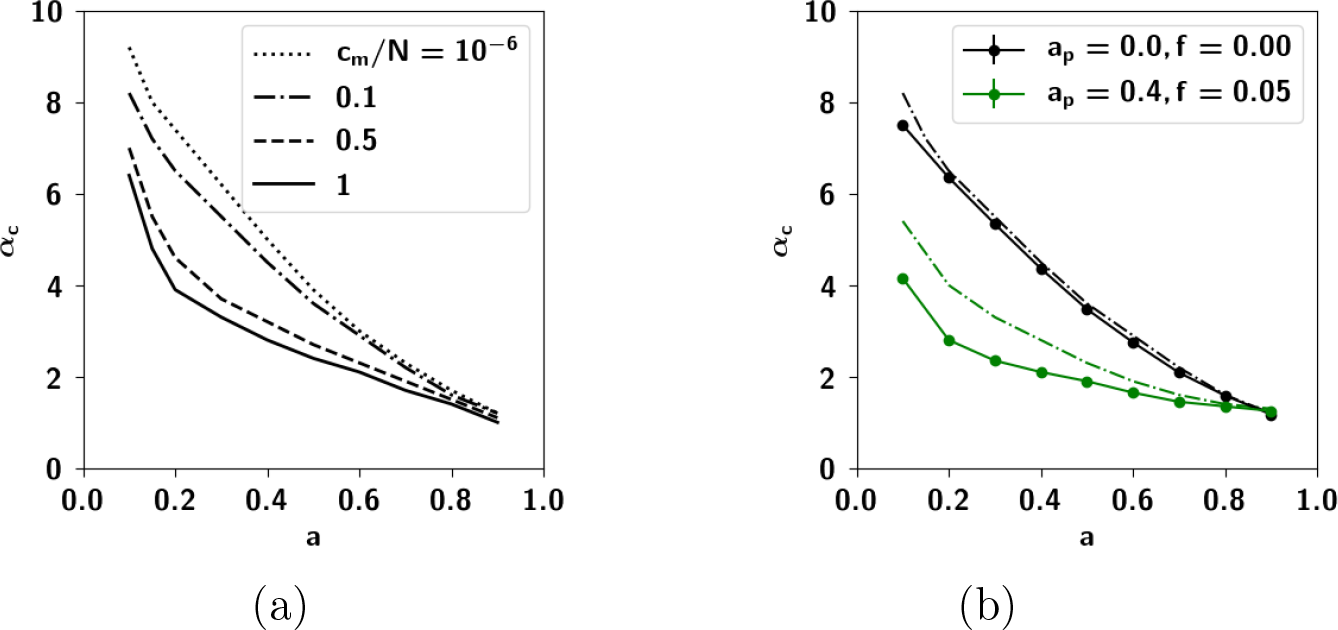
(**a**) Storage capacity as a function of the sparsity *a* for different values of the dilution parameter *c*_*m*_/*N*, for uncorrelated patterns 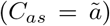 obtained through solutions of the mean-field equations. (**b**) Storage capacity for uncorrelated patterns (in black) and for correlated patterns (in green). Dots correspond to simulations while the dash-dotted lines to solutions of the mean-field equations. For uncorrelated patterns this is the same curve as in panel (a) with *c*_*m*_/*N* = 0.1. It is apparent that the mean-field treatment yields better results for uncorrelated patterns; for correlations, it over-estimates the storage capacity. Parameters are *N* = 2000, *c*_*m*_ = 200, *S* =5, *U* = 0.5 and *β* = 200.

In Fig. 16 we show the storage capacity for correlated patterns, for different values of the correlation parameters *a*_*p*_ and *f*. As can be seen in Fig. 16(a), increasing either extent of influence or prolificity, whatever the sparsity, is detrimental to the capacity. In Fig. 16(b), instead, we can see the capacity as a function of the number of Potts states *S*. For *S* = 1, as the algorithm produces uncorrelated patterns, the capacity remains the same, regardless of the correlation parameters. For higher values of S, on the other hand, the capacity decreases with increasing values of the correlation parameters *f* and *a*_*p*_. The behavior of the capacity as a function of *c*_*m*_, shown in the simulations of Fig. 16(c) which have been carried out with random dilution of the connectivity (see Eq. 7) shows the same strong dependence on correlations. The decrease in capacity brought on by the correlations is due to the increased variance of the noise, discussed in the previous section.

**Figure 16:**
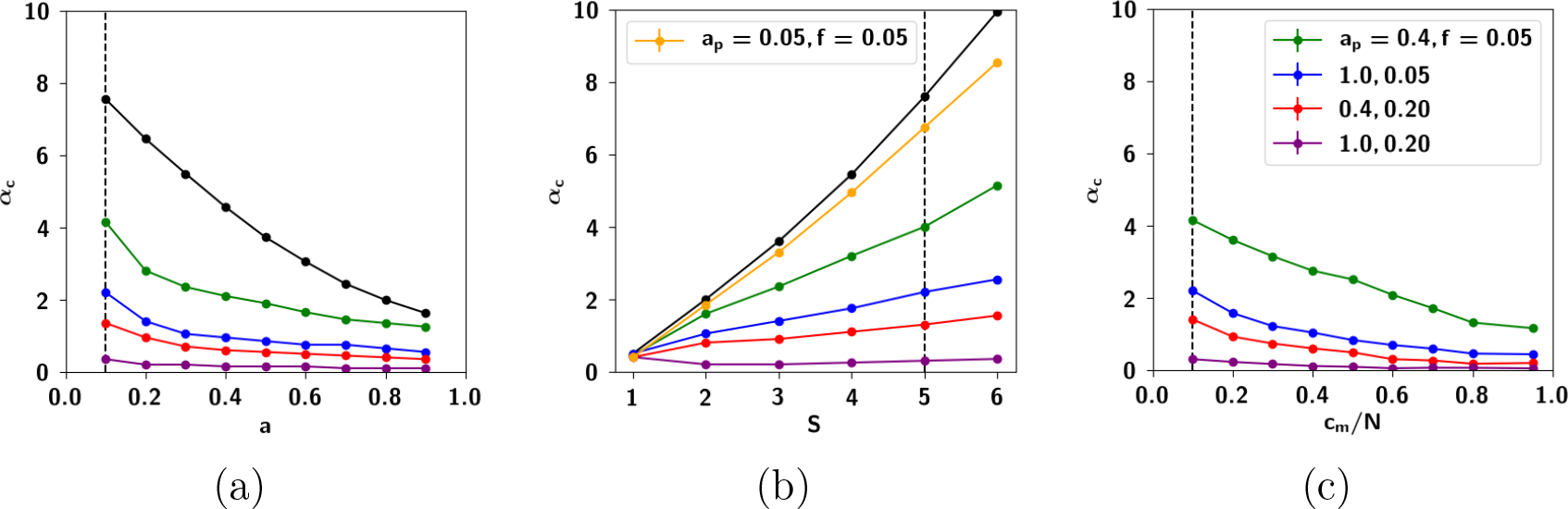
(**a**) Storage capacity *α*_*c*_ as a function of the sparsity *a* for different values of the correlation parameters *a*_*p*_ and *f*. The storage capacity is defined as the critical storage at which half of all cued patterns are retrieved with overlap of 0.7 and above. Increasing *a*_*p*_ and f are both generally detrimental to the capacity. (**b**) *α*_*c*_ as a function of the number of Potts states *S*, which shows that the superlinear increase derived analytically in [34] for randomly correlated patterns (the black curve) is only really approached, within this limited S range, with patterns that are very close to randomly correlated (the orange curve) (**c**) *α*_*c*_ as a function of the connectivity *c*_*m*_ for random dilution (see Eq. 7). The capacity decreases as a function of increasing connectivity. When not explicitly varied, parameters are *N* = 2000, *c*_*m*_ = 200, *a* = 0.1, *S* = 5, *U* = 0.5, *β* = 200, *ζ* = 10^−6^ and Π = 150.

#### 2.3.3 The effect of correlation parameters *f* and *a*_*p*_

In Fig. 17 we see the storage capacity as a function of the different correlation parameters *f* and *a*_*p*_. We can see that increasing each of these parameters decreases capacity, albeit in a different manner. The dependence of *a*_*c*_ on the prolificity f can be seen in Fig. 17(a): *a*_*c*_ decreases dramatically with increasing *f*, and goes to zero for very high values of *f*, in which children are each affected by a large number of parents. This result makes sense in light of the fact that *f* affects the mean correlation between children, as shown in Fig. 9(b).

**Figure 17:**
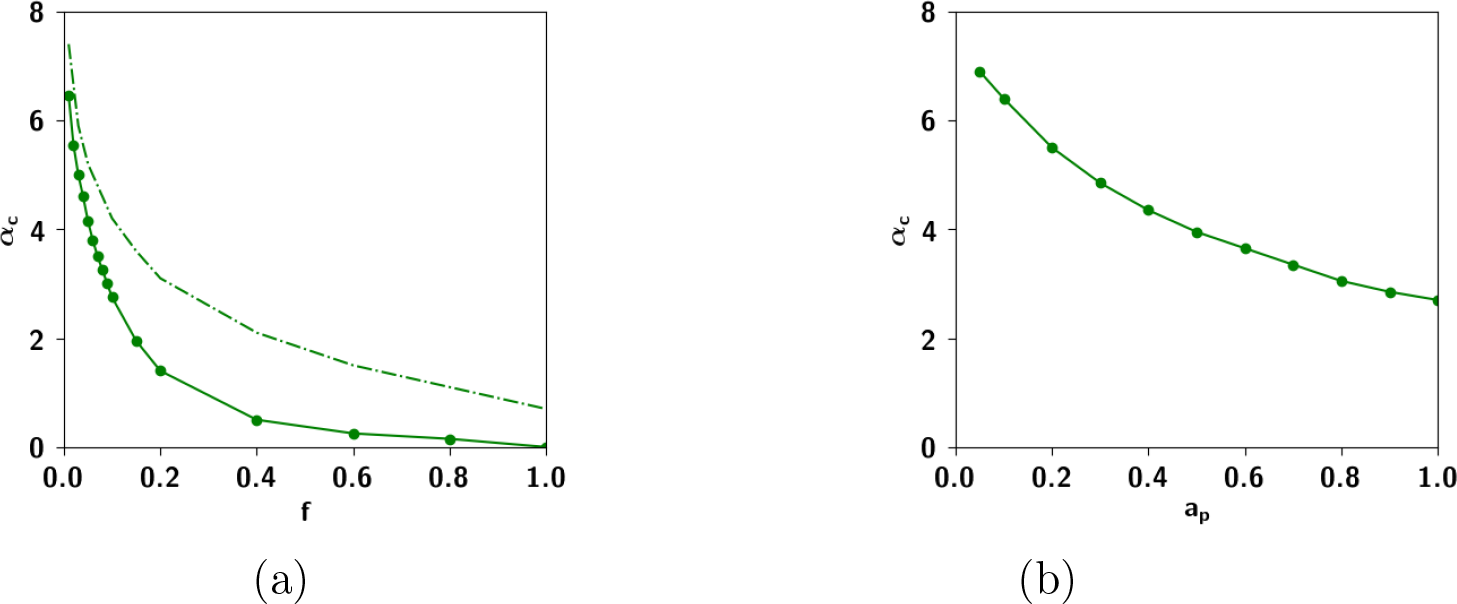
Storage capacity curves as a function of different correlation parameters. **(a)** *α*_*c*_ as a function of *f*. The full lines correspond to simulations while the dashed line corresponds to solutions of the mean-field equations. It can be seen that similar to Fig. 16(a), the over-estimation of the capacity through the SCSNA approach holds also for other values of *f*. **(b)** *α*_*c*_ as a function of *a*_*p*_. When not explicitly varied, the correlation parameters are *a*_*p*_ = 0.4, *f* = 0.05, *ζ* = 10^−6^ and Π = 150. Network parameters are *N* = 2000, *c*_*m*_ = 200, *a* = 0.1, *S* = 5.

In contrast, *α*_*c*_ decreases almost linearly with increasing the extent of parent input *a*_*p*_, as shown in Fig. 17(b), but does not go to zero for the highest possible value of *a*_*p*_ = 1. As we saw in the Sect. 2.2.5, *a*_*p*_ affects the degree to which children are similar to each of their individual parents. Increasing this parameter increases the similarity between those children receiving input from the same parents, increasing their overall similarity and therefore decreasing their discriminability. In terms of the effect on the correlation distribution, in Fig. 9(a) it can be seen that with increasing *a_p_,* there is an increase in the fluctuations in the correlations, making them more positively skewed.

#### 2.3.4 Correlated retrieval

In the previous section we saw that correlations decrease the storage capacity of the network. In particular, in terms of the dominance parameter *ζ*, what is the effect of correlations on memory retrieval? Which configurations of activity does the network settle into? We carried out simulations with correlated patterns for different values of *ζ*. We cued each pattern and, at the end of retrieval dynamics, we computed the overlap of the network configuration with all patterns and all parents. We then computed ⟨*m*_*cue*_⟩ as the mean overlap, over all simulations, of the network configuration with the cued pattern. Similarly we computed ⟨*m*_*corr*_⟩ as the mean overlap of the network configuration with another pattern, the highest among the children excluding the one cued. Finally we computed ⟨*m*_*fact*_⟩ as the mean overlap of the network configuration with a parent, the highest among all parents. We plot these values as a function of increasing loading *α* in Fig. 18.

**Figure 18:**
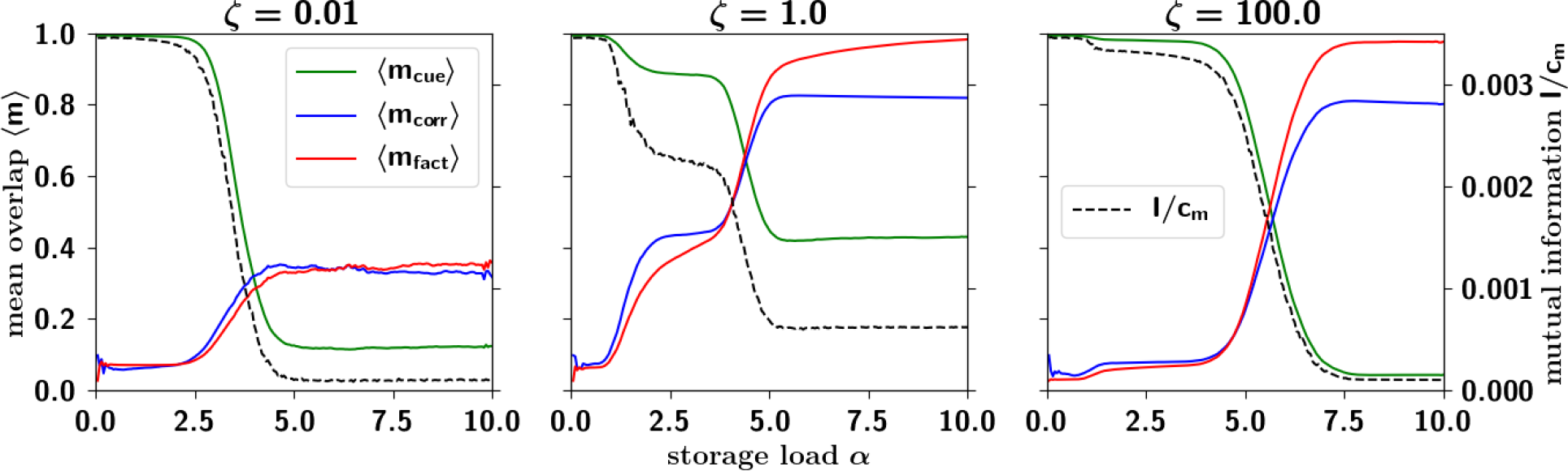
Mean overlap (left *y*-axis) and mutual information per connection (dashed black curve, and right *y*-axis) as a function of the storage load α, for different values of the dominance *ζ*. For low values of *ζ*, the information falls abruptly at a value of the storage load α, and similarly for very high values of *ζ*, while for intermediate values we observe a more gradual decay, starting at lower values of the storage load. For intermediate values of *ζ* however, the information does not go to zero, but rather saturates at a certain value. We call this *residual information*. In Fig. 19, we plot this residual information as a function of *ζ*. Network parameters are *N* = 2000, *c*_*m*_ = 200, *S* = 5, *a* = 0.1, *U* = 0.5, *β* = 200. Correlation parameters are *a*_*p*_ = 0.4, *f* = 0.05 and Π = 150.

We can see that for low values of *ζ*, the fall of ⟨*m*_*cue*_⟩ is accompanied by only a modest increase in both ⟨*m*_*corr*_⟩ and ⟨*m*_*fact*_⟩, that we call respectively *correlated* and *factor retrieval*. Increasing *ζ*, here by two orders of magnitude, we observe two stages: a first (partial) fall in ⟨*m*_*cue*_⟩, accompanied with a moderate increase in ⟨*m*_*corr*_⟩ and ⟨*m*_*fact*_⟩; and then a second fall, after which ⟨*m*_*corr*_⟩ and ⟨*m*_*fact*_⟩ exceed the former. Increasing *ζ* by another two orders of magnitude, cued retrieval is restored to more than its initial value (with *ζ* small) beyond which we observe *factor* retrieval.

This effect is summarized in Fig. 19, the storage capacity *α*_*c*_ as a function of *ζ*, where it can be seen that the capacity displays a trough at intermediate values of the dominance *ζ*. At lower values of *ζ*, well below the trough, the basins of attraction are rather well-separated (as we can anticipate from the clustering analysis of the next section, summarized in Fig. 20(a)). The large barriers mean that each individual pattern can be retrieved relatively well.

**Figure 19:**
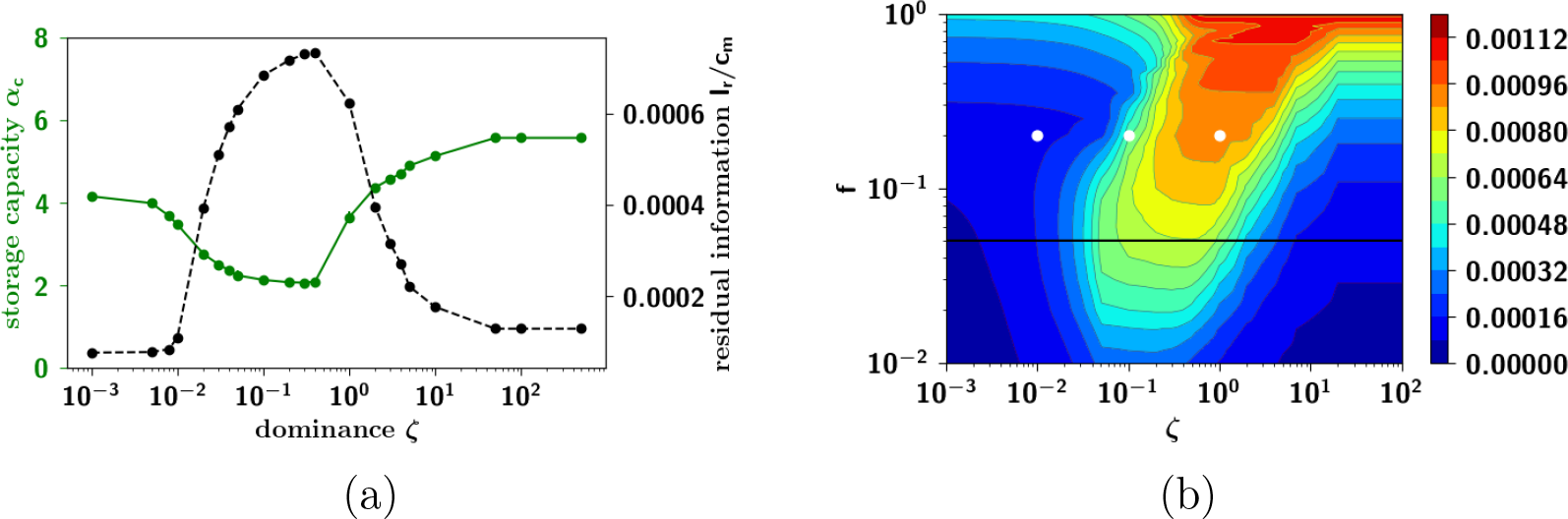
(**a**) Storage capacity (left *y*-axis) and residual mutual information between cued memory and configuration retrieved, after capacity collapse, as a function of *ζ* (right *y*-axis). The storage capacity displays a trough for intermediate values of the dominance *ζ* that is due to an increased clustering of the patterns and the inability of the network to retrieve each one of them with relatively high precision. The apparent increase of the capacity, for very high values of the parameter *ζ*, is instead due to such clusters vanishing altogether, one by one, as the inputs from weaker parents are dominated by small input to a random state (see Eq. 31). The residual mutual information corroborates the results from the storage capacity: it is only for intermediate values of *ζ* that the network can retrieve some information after capacity collapse. Network parameters are *N* = 2000, *c*_*m*_ = 200, *S* = 5, *a* = 0.1, *U* = 0.5, *β* = 200. Correlation parameters are *a*_*p*_ = 0.4, *f* = 0.05 and Π = 150. (**b**) Phase diagram of the residual information, as a function of the dominance *ζ* in the *x*-axis and prolificity f in the *y*-axis, giving a fuller picture of the phase transition of Fig. 19(b). Note that the transition to non-zero residual mutual information occurs at higher values of *ζ* with increasing *f*. The black horizontal line, plotted for clarity, corresponds to the value of *f* used in Fig. 19(a). The three white dots correspond to three points for which a cluster analysis is reported in Fig. 20.

**Figure 20:**
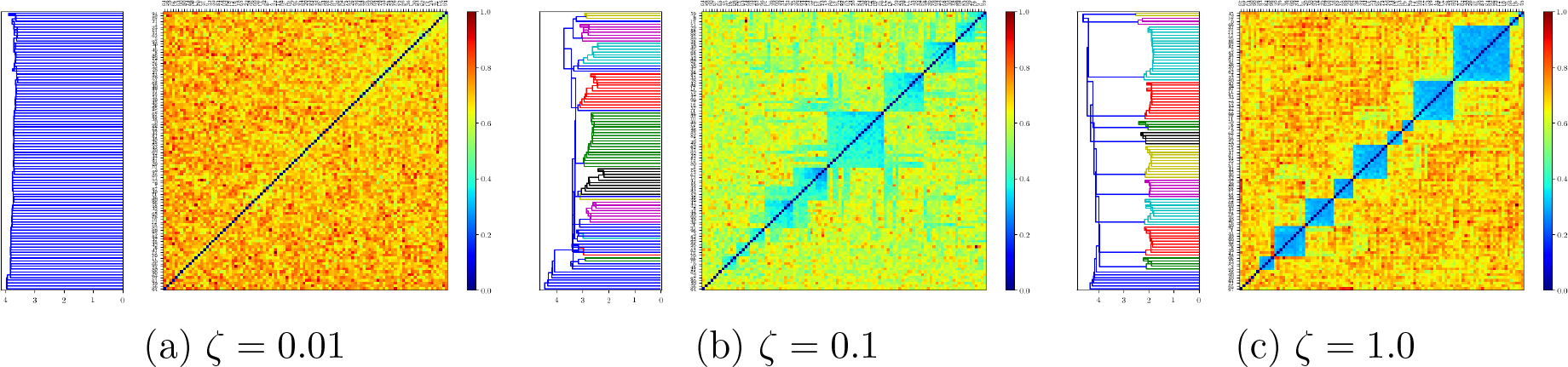
Cluster analysis applied to patterns generated by the algorithm, for three different values of the dominance parameter *ζ*, chosen at salient points of the phase diagram in Fig. 19. Increasing *ζ*, the patterns generated by the algorithm become more and more clustered, as the strongest parent of each pattern comes to dominate its activity.

Increasing *ζ*, up to where the capacity reaches its minimum value, patterns start to become increasingly clustered, and the barriers between them smaller. In this regime of intermediate clustering, the network can get confused during retrieval, effectively stabilizing into another stable state. This decrease in the capacity is however accompanied by correlated and to a greater extent factor retrieval: the network is not able to distinguish between individual patterns but recognizes the cluster it belongs to. Such a behavior has been found in previous models exploring correlations, and in particular in [44], where an ultrametric organization of the patterns is not a limit outcome of the model as in ours (see Fig. 20) but is the object of study itself. In the latter study, the temperature is the control parameter, below a certain critical value of which the network can distinguish individual patterns, and above which the network can recall clusters, but not the individual patterns within each cluster.

As *ζ* is increased even further, the capacity starts to increase, eventually saturating at a given constant value. This increase of the capacity is due to the elimination of clusters, as the input from the weaker parents become comparable to the input from a randomly chosen state e in Eq. 31. Those clusters of patterns having as strongest parent, one of the weaker parents (in absolute) become eliminated one by one until there is only a single cluster left, corresponding to patterns belonging to the first (and strongest) parent. Moreover, that the storage capacity at such high values of *ζ* is higher than that at very low values of *ζ* is probably due to the fact that at high *ζ*, those patterns not belonging to the single cluster are randomly correlated with one another, while patterns at very low *ζ* are weakly correlated with one another. Such high values of *ζ* are, however, hardly plausible in terms of semantic organization, and hence outside the scope of our interest.

#### 2.3.5 Residual information: memory beyond capacity

We can further corroborate the findings in the previous section through the mutual information between the pattern cued *c* and the configuration in which the network settles *r* 
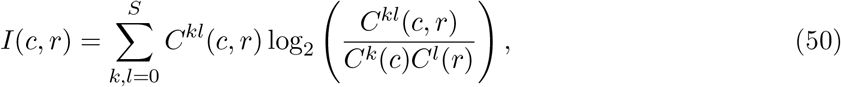
where 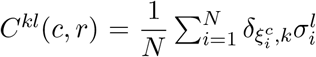,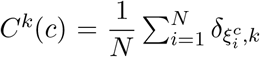 and 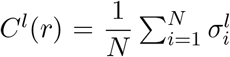. The maximum value of this quantity is attained when the cued pattern is also the one retrieved: *c* = *r*. In this case the mutual information can reach up to 
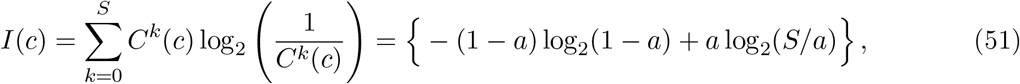
that we recognize to be the entropy of the cued pattern. In Fig. 18 we can see the mutual information as a function of the loading α for different values of the parameter *ζ*, averaged across cued retrieval of many patterns. For small values of α, the mutual information does not depend on *ζ*: its value at this plateau corresponds to the entropy. For small *ζ*, the mutual information has a sharp fall-off upon increasing *α*, falling to approximately zero. For the intermediate value of *ζ* reported, it displays a step-like behavior, but ultimately stabilizes to a constant non-zero value. Increasing *ζ* even further, it again has a fall-off, at a higher value of the storage load, qualitatively following the behavior of the overlap in the same figure.

The most interesting observation in Fig. 18, however, is the *residual information*, its remaining roughly constant value, after capacity collapse, in a range of intermediate values of *ζ*. In Fig. 19(a), this residual information is plotted as a function of the dominance rate *ζ*, and it can be seen that it increases sharply at approximately *ζ* ≃ 0.01 (reaching a value, given these parameters, of order 0. 7 × 10^−3^ bits per connection, some five times below the entropy of the stored pattern, the initial plateau in Fig. 18). It then decreases again at approximately *ζ* ≃ 0.5. This effect is reminiscent of a phase transition with control parameter *ζ*, where the information plays the role of the order parameter. Below *ζ* ≃ 0.01 and above *ζ* ≃ 0.5, once the capacity is exceeded, there is no more retrievable information about the cued pattern. Within the range 0.01 ≳ *ζ* ≳ 0.5 however, the network retrieves some information about the cued pattern. In Fig. 19(b) we plot, as a phase diagram, the residual information as a function of *ζ* in the *x*-axis and *f* in the *y*-axis. One sees that a non-vanishing residual information requires, essentially, intermediate values of *ζ* and sufficiently large values of *f*. In terms of either parameter, the region with non-vanishing residual information spans more than one order of magnitude. This intermediate regime can be argued to form the basis of semantic resilience in our model.

#### 2.3.6 Residual memory interpreted through cluster analysis

Although it was argued in the introduction that the presence of clusters is only one component of the metric relations embedding semantic memories, it is nevertheless instructive to interpret the residual memory phase expressed by our model in terms of cluster analysis. Fig. 20 shows the outcome of applying a standard clustering algorithm to patterns generated at salient locations of the phase diagram in Fig. 19(b). For low dominance (*f* = 0.2 and *ζ* = 0.001, Fig. 20(a), i.e. weak correlations), the algorithm is essentially unable to identify clusters: all patterns seem to be at roughly equal distance from each other. For higher dominance (*ζ* ≃ 0.1) the clustering structure emerges, as indicated by the expanding white area to the left. In the high dominance region (*ζ* ≃ 1, i. e. strong correlations), a few parents dominate the rest and the extracted clusters are clearer, accompanied with a concomitant increase in the residual information. For even larger *ζ* values there is only one cluster (not shown here), effectively, and no residual information above *α*_*c*_, which returns to a value above that for weak correlation – as they are again effectively random, against the backdrop of that single parent, which acts as a biased probability distribution. We can conclude, therefore, that the residual information largely expresses resilient memory associated with the distinction between clusters, whereas within-cluster distinctions are lost above the capacity limit.

An interpretation of these results can be framed in terms of *categorical perception* [45]. Whenever we learn categories, it becomes more difficult to distinguish two patterns within a category than two patterns straddling the boundary between two categories, even when the two pairs are at the same physical distance. Within-category discrimination is reduced while between-category discrimination is enhanced. In addition to this, there exists a perceptual warping within the category. All categories acquire an internal structure: how well each pattern fits in the category can be measured with a *goodness* metric [46, 47] and the element with the largest goodness is called the prototype. Whenever a pattern is observed, it is perceived as closer to the prototype of the category than it really is. This effect is called the *perceptual magnet* [48] and has been observed in many contexts, see [49] for a review.

The results we obtain here, although pertaining to a model of semantic memory not of perception, and a model that transcends the simple-minded notion of category, are reminiscent of these phenomena. For moderate values of *ζ*, the exact cued pattern may be lost but still the final pattern belongs with it in some cluster. The fact that the final pattern is correlated to the strong parent pattern is akin to the perceptual magnet if we think of the parent pattern as embodying the cluster, while the child patterns are distortions of it. We recently gave an account of perceptual phenomena using as central feature the similarity between patterns [15] and showed that the existence of the structure of a category and the perceptual effects are natural consequences of the similarity structure of the patterns within a category. In both approaches, taking into account the similarity (or correlation) structure of a set of patterns lead, in some regimes, to a similar phenomenology. However, this connection remains qualitative, in that our pattern generation algorithm does not produce well-defined categories, but a more complex set of metric relations among patterns, where clusters emerge as one component if a few parents dominate.

#### 2.3.7 Residual memory rides on fine differences in ultrametric content

A complementary perspective is that afforded by our measure of ultrametric content, which is derived from a measure of distance between patterns, applied to all triplets of patterns. A suitable distance measure, for Potts patterns, can be 
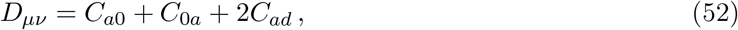
where similar to the quantity we introduced in Sect. 2.2 (*C*_*as*_), *C*_*ad*_ is the fraction of co-active units which are in different states in both patterns *μ* and *ν*. *C*_0*a*_ is the number of units quiescent in *μ* and active in *ν* and conversely *C*_*a*0_ is the number of units active in *μ* and quiescent in *ν*. In the dominance-prolificity region we have been considering, this distance measure yields the values seen in Table 1 for the ultrametric content index (see App. C).

**Table 1:** Ultrametric content computed for distances of triplets of patterns generated by the algorithm, for six different parameter values of the prolificity and the dominance. An increased ultrametric content reflects an increased clustering in the correlations between patterns. For *f* = 0.2 and *ζ* = 0.1, the patterns yield values of the ultrametric content index close to that obtained from the nouns (~ 0.5). The corresponding clustering structure of the patterns can be seen in Fig. 20(d).

| | | $\zeta$ | | | | |
| --- | --- | --- | --- | --- | --- | --- |
|  |  | 0.001 | 0.02 | 0.1 | 1 | 10 |
| $f$ | 0.05 | 0.395 | 0.429 | 0.435 | 0.416 | 0.397 |
|  | 0.2 | 0.389 | 0.404 | 0.507 | 0.507 | 0.507 |

Interestingly, a completely different measure of distance, similar instead to the one extracted from the feature-based norms in the toy example reported in Sect. 1.1, yields values very close to these, within a few percent, when applied to patterns generated with our algorithm. We can see, therefore, that the emergence of residual memory does correlate with increased ultra-metric content, but not in a simple one-to-one correspondence; and the putative phase transition to non-zero residual information occurs in the midst of a relatively minor increase in the values of the ultrametric content index.

A tentative conclusion is that semantic resilience, at least as crudely modelled by a Potts network, requires a degree of clustering or ultrametric structure, which in the pattern-generation model reflects sufficient values of prolificity and dominance, but is still an emergent phenomenon. Quantitative differences in the parameters produce what tends to be an all-or-none difference in semantic resilience. Yet another form, possibly, of analog-to-digital transform produced by a neural circuit.

## 3 Discussion: a new model for the extraction of semantic structure

In recent years, the Potts network has been proposed as an effective model of a cortical network organized with distinct local and global connections. Several aspects of the model have been studied in quantitative detail, under many simplifying assumptions, including that of uncorrelated patterns. However, it can be argued that such patterns are irrelevant for the study of semantic memory. The various feats of “mind-reading”, achieved with fMRI studies (e.g. [50, 51, 7]), reflect that correlations between memories are not simply a nuisance that degrades memory capacity, but express the core ability of the cortex as a machine for encoding structured information. We have attempted to make theoretical progress in this direction by designing a plausible algorithm to generate patterns. In our algorithm, the patterns are generated by the contribution of multiple factors, which can be considered as semantic category generators (except that categories overlap and have loose boundaries), or else features in a somewhat wider sense, that carry information on the statistical co-occurrence of attributes. Through competition, those attributes concur, each with its relative strength (parametrized by *a*_*p*_, *f* and *ζ*) to construct the statistical structure of the memories.

We have further studied the storage capacity of the network as a function of both network parameters and correlation parameters through the SCSNA analysis as well as extensive simulations. We find that with a Hebbian rule for the storage of patterns, the network can store and retrieve *fewer* correlated patterns, though still of order *c*_*m*_*S*^2^/*a* (with weakly correlated patterns), yielding ~ 10^7^ with human cortical parameters [31]. Other prescriptions for learning, enhancing capacity, may be explored and studied, and we leave such studies for future investigations. Of the correlation parameters, the effect of the dominance *ζ* is particularly interesting. *ζ* ≃ 0 corresponds to a situation in which all parents are on equal footing, while the opposite limit corresponds to only one handful dominating the rest. For intermediate values of the dominance *ζ*, we observe *correlated retrieval*, in that with the decrease in successful retrieval, the fraction of trials in which another, correlated pattern is retrieved, increases; a closer look however, indicates that the phenomenon is linked to the retrieval of the factors, i.e. *factor retrieval*. In terms of the mutual information between the cued pattern and the final configuration of the network, after retrieval dynamics, we observe that in an intermediate regime of *ζ*, after the capacity limit has been reached, it does not go to zero. We call this the *residual information*. The residual information displays a non-monotonic dependence on the dominance *ζ*. Such a nontrivial behavior is reminiscent of a phase transition, in which the residual information is the order parameter and *ζ* is the control parameter. The residual information has an interesting interpretation: it can be thought of as the information pertaining to the *gross*, core semantic component of the memories, after the *fine* details have been compromised. Note that 1/*ζ* is a measure of the number of parents/factors/attributes that effectively dominate semantic space.

Taken at face value, the diminished capacity of the Potts network accompanied by the emergence of residual information suggests that this ability for generalization comes at the cost of losing the resolution with which we can retrieve the individual memories. However, this result as such is incomplete, and must be considered also in relation to the differential role of other memory structures and in particular the hippocampus in retrieval. For example, in humans, it has been shown that the ventral hippocampus projects directly to the medial prefrontal cortex, providing an immediate route for representations to reach the prefrontal cortex, suggesting a model of bidirectional hippocampus/prefrontal cortex interactions that support context-dependent memory retrieval [52]. Several studies have attempted to dissociate between the contributions of the hippocampus and the cortex in human memory retrieval. In particular, in [4] and [53] it was found that putatively different access modes to information stored in long-term memory in a remember/know paradigm lead to different distributions of classification errors of different groups with memory disorders. An information derived measure, the metric content, quantifying the concentration of errors was computed: high levels of metric content are indicative of a strong dependence on perceived relations among the set of stimuli, and therefore of a relatively preferred semantic access mode, while low levels (and similar correct performance), suggest a preferential episodic access mode. It was found that compared with normal controls, the metric content index was increased in patients with Alzheimer’s disease, decreased in patients with herpes encephalitis, and unvaried in patients with damage to the prefrontal cortex. Moreover, a significant correlation between the metric content and measures quantifying episodic and semantic retrieval mode in the remember/know paradigm introduced by Tulving [54] was found. If we think of the access modes, to a first approximation, as reflecting a stronger reliance on specific memory structures, the distribution of errors may then be a window into understanding their relative contributions. Within this larger picture, a cortical impairment as modelled for example by a Potts network with reduced connectivity may be somewhat mitigated by a complementary episodic mode of access, supported by other structures.

Our finding of semantic resilience, as characterized by the residual information has an interesting interpretation also in relation to the findings in the neuropsychology of semantic dementia. A typical finding is that the finer-grained or “subordinate” aspects of such patients’ knowledge are more susceptible to damage than the more “ordinate” aspects, for example in naming tasks. Moreover, the naming errors that such patients make tend to change in time from “circumlocutions to category coordinates to superordinate labels” [55]. It has been argued that such a finding is in favor of treelike models of semantic knowledge, in which the mental representation of a concept occupies nodes in a branching tree, where the origin of the tree corresponds to its most general and the periphery to its most selective designation. Subsequent research however, pointed to findings that could not be explained through a tree model, such as verification latency [56] or typicality effects [57], or others which question where in the tree to store concepts that belong to more than one category [58]. In our account, instead, semantic resilience is an outcome that emerges naturally through higher values of the dominance parameter *ζ*, in which finer-grained or subordinate features of the concepts are overtaken by ordinate features, which then become the only retrievable ones, as shown through the cluster analysis in Fig. 20. Crucially though, such clusters emerge only when our dominance parameter becomes large enough, and they are neither well-defined nor designed to have strict boundaries. The non-trivial behavior of the residual information, i.e. its phase transition with the dominance parameter *ζ*, cannot be predicted from a qualitative model encoding uncorrelated patterns. Our model offers one plausible way in which such resilience emerges.

Finally, our account may have implications for the question of how the cortex extracts and encodes the general statistical structure of the ensemble of stimuli that it receives. There is a well-established view of the cortex as a slow memory system that uses overlapping distributed representations to represent the general statistical structure of the environment. It has been suggested that the interaction between the hippocampus and the cortex is a crucial element in the consolidation of memories. The general idea is that memories are first stored in the hippocampal system via synaptic changes and that these support the reinstatement of recent memories in the neocortex. Neocortical synapses are slightly modified on each reinstatement and the gradual, neocortical changes accumulating over time encode remote memory. This division of labor would allow the hippocampus to rapidly encode new episodic items without disrupting semantic memories, and the cortex to slowly integrate them in a structured fashion into such memories.

This view is consistent with evidence that damage to the hippocampal system results in recent memory disruption but leaves remote memory intact, but it does not really specify what makes the consolidation process gradual or slow [59]. Early modelling attempts typically resorted to backpropagation to account for the structured learning of the cortex [60]: consolidation would then be slow, because backpropagation is effective with low learning rates. However, backpropagation has been widely criticized on the basis that it lacks a plausible biological mechanism. While hippocampal learning in these accounts was taken to fit the framework of learning unrelated patterns of activity, it had remained unclear how to model neocortical learning. Our account offers an alternative framework for neocortical learning, in which semantic structure is extracted progressively from the statistics of generating features and encoded in the cortex via Hebbian learning and is resilient, i.e. it is preserved when the storage capacity for “episodic details” is exceeded.

### A Calculation of the probability distribution of the field for *S* = *1*

In this section we outline the main steps in deriving Eq. (25). The distribution for *h*_μ_ can be computed by making use of the probability generating function 
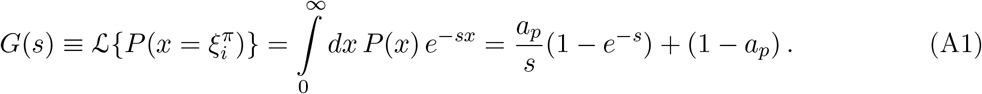

Since the 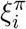 are identically and independently distributed for all *π*, we can use the following property 
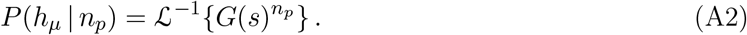

The number of parents as well as the total field received by a pattern is i.i.d, so we drop the index *μ* for *h*_μ_. We can compute the conditional distribution of the field received by a unit, given the number of parents, as 
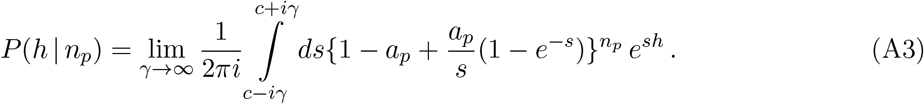

Using the binomial theorem 
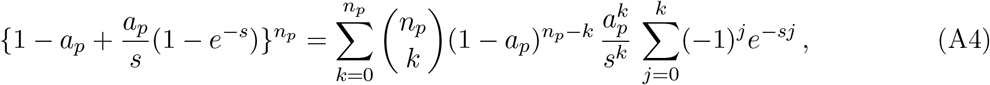

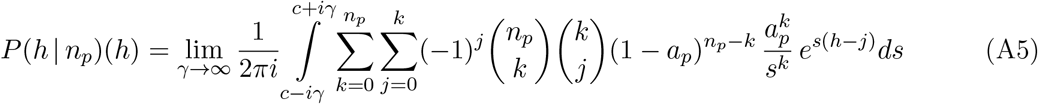

We can carry out the integral to find 
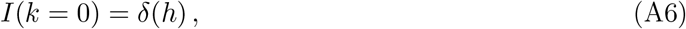

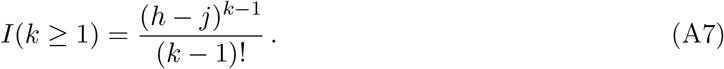

The distribution of the field *h* for a given number of parents *n*_*p*_, is then 
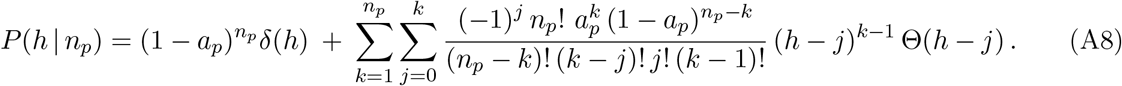

The first term in this equation expresses the fact that the only way to get zero field is if all *n*_*p*_ parents contribute zero field and this occurs with probability (1 – *a*_*P*_)^*np*^. For a given pattern *μ*, with *n*_*p*_ parents, the field of each unit is distributed according to Fig. 6. The cumulative distribution function writes 
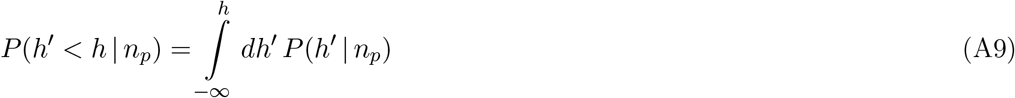

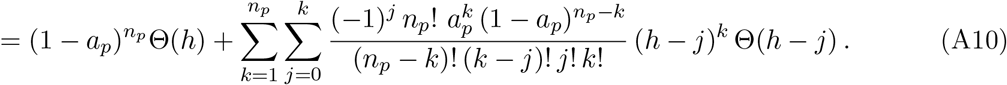

The minimal threshold *h*_*m*_ is implicitly given by the cumulative probability 
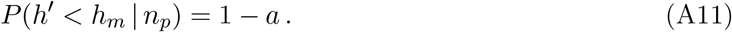

### B Calculation of the probability distribution of the field for *S* = *2*

To derive Eq. (29), we start with the joint distribution of number of parents by state 
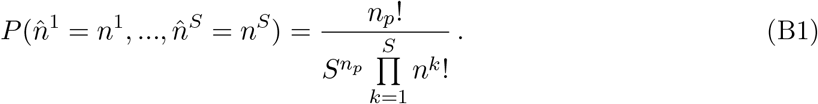

Note that we define the field to be identically distributed across states. The probability that the fields of all states are below that of the first is given by 
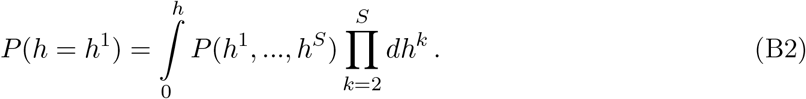

The probability distribution of the maximal field is given by *S* times the one above 
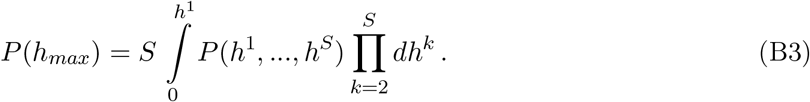

The joint distribution of the fields across states writes 
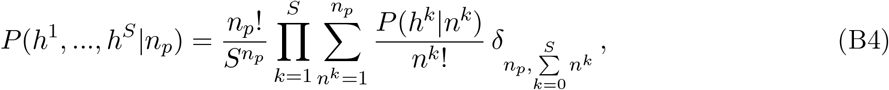
where the constraint 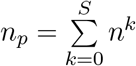 has been included in the last line. *P*(*h*^*k*^ ~*n*^*k*^) is given by Eq. (A8), replacing *n*_*p*_ with *n*^*k*^. We then have 
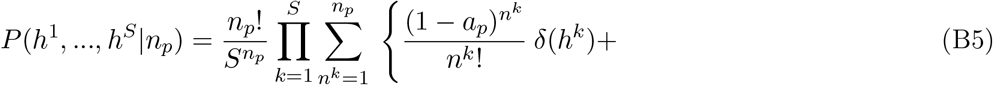

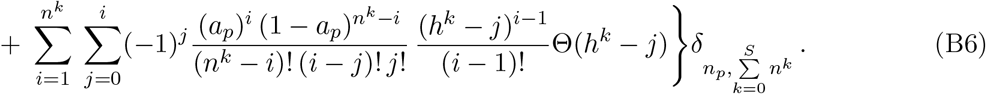

For *S* = 1 all contributions go to a single state, so we automatically have *n*^1^ = *n*_*p*_, then the first sum disappears and we fall back onto Eq. (A8). For *S* = 2 we have, denoting the state receiving the maximal field by *H*, 
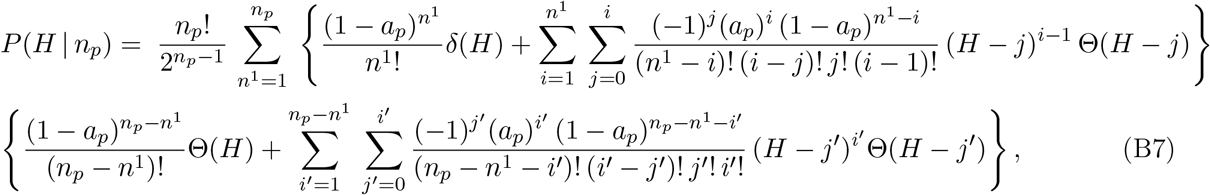
where we drop the indices denoting the units (they are drawn from the same distribution). Note that the state does not appear in this expression because it is the distribution for the state that receives maximal input, regardless of which one it is. The *μ* dependence is through *n*_*p*_ = *n*_*p*_(*μ*). We then get the minimal threshold for activation *H*_*m*_ implicitly in terms of the cumulative distribution 
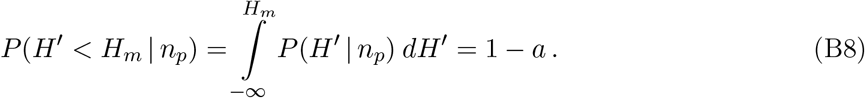

We can compute it to find 
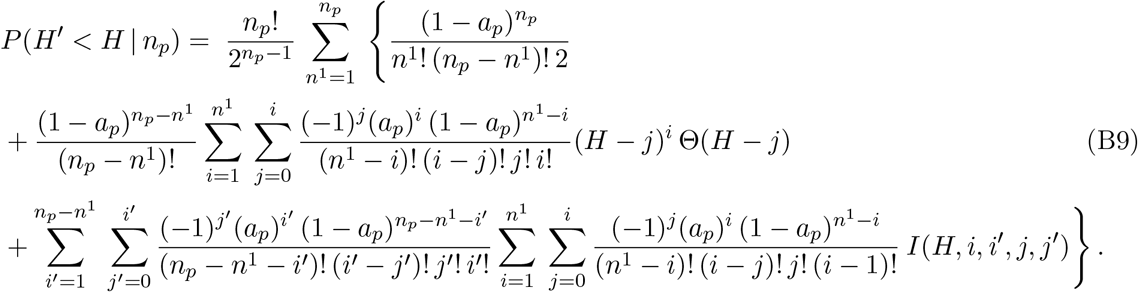
where *max*{*j*, *j′*} = *j*\* 
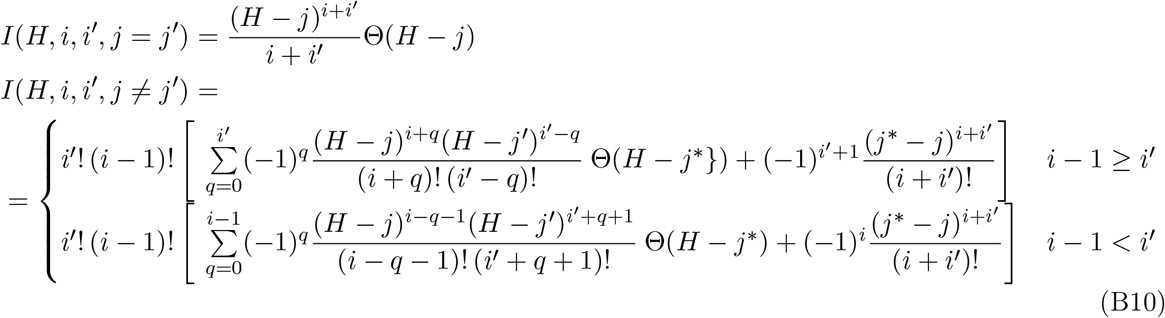

### C Ultrametric content

A possible characterization of the correlations between the memory patterns is in terms of a distance. A quasi-distance measure can be derived from the correlation following the same procedure as in [10]. We first define a so-called “confusion” matrix 
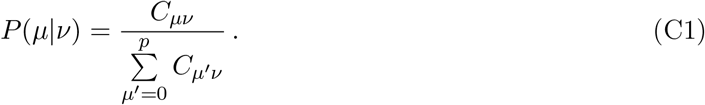
where *C*_*μν*_ is an element of the correlation matrix and where *P*, the confusion matrix, is obtained by normalizing each element of the correlation matrix appropriately. Next, we symmetrize the above function to obtain 
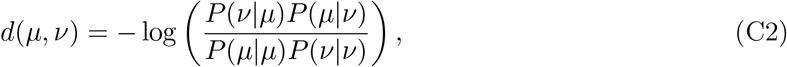
a quasi-distance, in the sense that it satisfies only the reflective and symmetric properties, *d*(*μ, μ*) = 0 and *d*(*μ, ν*) = *d*(*ν, μ*). The triangular inequality *d*(*μ, ν*) + *d*(*μ, ρ*) ≤ *d*(*μ, ρ*) does not necessarily hold. It can be made to hold by raising d to a sufficiently small power *d* ↔ *d*^1/*p*^, called the “trivialization” of *d*, as explained in detail in [10]. Using this procedure, distances between triplets of patterns {*μ, ν, ρ*} can be computed. If we note by *d*_*min*_ the edge of minimal length, *d*_*max*_ the edge of maximal length and *d*_*med*_ the edge of intermediate length, then we can plot, in a two-dimensional graph, the ratios *δ*_1_ = *d*_*min*_/*d*_*max*_ and *δ*_2_ = *d*_*med*_/*d*_*max*_. Triplets that satisfy the triangular inequality lie above the line *δ*_1_ = 1 – *δ*_2_, while triplets that satisfy the ultrametric inequality lie on the vertical line where *δ*_2_ = 1. Among these, triplets that are equilateral triangles lie at the point *δ*_1_ = *δ*_2_ = 1. To measure the overall closeness of the cloud of triplets to the fully ultrametric limit one can define the *ultrametric content* 
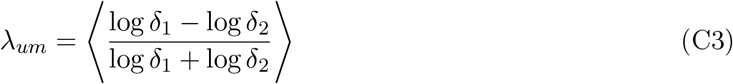
where ⟨·⟩ denotes the mean over all triplets. This quantity does not depend on the trivialization of d and it ranges from 0 (for triplets forming isosceles triangles with two short sides) to 1 (for a fully ultrametric set: equilateral triangles and isosceles triangles with two long sides).

